# Microglial phagocytosis dysfunction during stroke is prevented by rapamycin

**DOI:** 10.1101/2021.11.12.468358

**Authors:** S Beccari, V Sierra-Torre, J Valero, M García-Zaballa, A Carretero-Guillen, E Capetillo-Zarate, M Domercq, PR Huguet, D Ramonet, A Osman, W Han, C Dominguez, TE Faust, O Touzani, P Boya, D Schafer, G Mariño, E Canet-Soulas, K Blomgren, A Plaza-Zabala, A Sierra

## Abstract

Microglial phagocytosis is rapidly emerging as a therapeutic target in neurodegenerative and neurological disorders. An efficient removal of cellular debris is necessary to prevent buildup damage of neighbor neurons and the development of an inflammatory response. As the brain professional phagocytes, microglia are equipped with an array of mechanisms that enable them to recognize and degrade several types of cargo, including neurons undergoing apoptotic cell death. While microglia are very competent phagocytes of apoptotic cells under physiological conditions, here we report their dysfunction in mouse and monkey (*Macaca fascicularis* and *Callithrix jacchus*) models of stroke by transient occlusion of the medial cerebral artery (tMCAo). The impairment of both engulfment and degradation was related to energy depletion triggered by oxygen and nutrients deprivation (OND), which led to reduced process motility, lysosomal depletion, and the induction of a protective autophagy response in microglia. Basal autophagy, which is in charge of removing and recycling intracellular elements, was critical to maintain microglial physiology, including survival and phagocytosis, as we determined both in vivo and in vitro using knock-out models of autophagy genes and the autophagy inhibitor MRT68921. Notably, the autophagy inducer rapamycin partially prevented the phagocytosis impairment induced by tMCAo in vivo but not by OND in vitro. These results suggest a more complex role of microglia in stroke than previously acknowledged, classically related to the inflammatory response. In contrast, here we demonstrate the impairment of apoptotic cell phagocytosis, a microglial function critical for brain recovery. We propose that phagocytosis is a therapeutic target yet to be explored and provide evidence that it can be modulated in vivo using rapamycin, setting the stage for future therapies for stroke patients.

## INTRODUCTION

Stroke is one of the most pervasive neurological diseases, with a global estimated yearly incidence of 13.7 million people that suffer disabling motor and neurophysiological deficits. It is also one of the most lethal diseases, claiming 5.5 million lives every year ^1^. The pathophysiology of stroke is particularly complex, with a cascade of events initiated by the lack of blood supply resulting from a broken or blocked blood vessel. A key element throughout this process is the immune system and particularly the macrophages residing in the brain parenchyma, microglia ^2^.

The dual role of microglia as a double-edged sword orchestrator of the brain immune response is well-recognized ^2, 3^. This dichotomy is evident in the contradictory results obtained by different studies using microglial depletion paradigms in rodent stroke models, with either detrimental ^4 5^ or beneficial ^6^ consequences. On one hand, microglia control inflammation and infiltration of peripheral immune cells, with deleterious effects on neurons when sustained or uncontrolled. On the other hand, microglia have beneficial effects by engulfing monocytes ^7^, neutrophils ^8^, and damaged blood vessels ^9^, controlling astrocytic inflammatory cytokines ^5^, or releasing growth factors such as IGF1 ^4^. In spite of these examples, functional studies of microglia in stroke remain scarce.

One indispensable microglial function that has received little attention is the phagocytosis of cellular debris generated during stroke. Efficient phagocytosis is crucial for the recovery of the homeostasis of the brain parenchyma, not only because it removes apoptotic corpses before they progress into secondary necrosis, releasing toxic intracellular compounds; but also because it modulates the phagocytés inflammatory response ^10^. We do know that phagocytosis is beneficial in stroke because its genetic ^11^ or pharmacological ^12, 13^ inhibition results in larger infarct areas. However, we still lack a basic understanding of how successful phagocytosis during stroke is.

The high efficiency of microglial phagocytosis of apoptotic cells under physiological conditions ^14^ is in sharp contrast to the dysfunction we recently reported in mouse and human epilepsy ^15, 16^. Epilepsy and stroke share some cardinal neuropathological events such as inflammation and excitotoxicity, thus raising the possibility of microglial phagocytosis impairment during stroke. To test this hypothesis, we used in vivo models of transient medial cerebral artery occlusion (tMCAo) and hypoxia-ischemia in mice and monkeys, as well as in vitro models of oxygen and nutrient deprivation (OND) in organotypic cultures and primary microglia. We found a pervasive impairment of microglial phagocytosis of apoptotic cells in all models studied that was related to several cellular mechanisms, including the induction of a protective autophagic response due to the energy depletion associated with stroke. We found that inhibition of basal autophagy using genetic and pharmacological approaches was essential to maintain microglial survival and function. In contrast, inducing autophagy with rapamycin did not improve the OND-induced phagocytosis blockade and had a deleterious effect on phagocytic microglia in culture. Nonetheless, rapamycin prevented to some extent the impairment of phagocytosis induced by tMCAo, supporting the possibility of pharmacological modulation of microglial phagocytosis in vivo.

## RESULTS

### Phagocytosis impairment in mouse and monkey models of stroke

We studied the impact of stroke on microglial phagocytosis in mouse and monkey models of transient occlusion of the medial cerebral artery (tMCAo; **Figure 1**). In mice we focused on the hippocampus, where we can establish the baseline of microglial phagocytosis efficiency in control conditions because of the ongoing apoptosis of neural progenitors in the hippocampal neurogenic niche ^14^, allowing us to quantitatively compare it with disease models ^15^. To ensure that the hippocampus was systematically affected by the MCAo we used an extended tip filament that blocked the collateral branches of the MCA, including the anterior choroidal artery, which irrigates the hippocampus (**Figure 1A**). We used laser Doppler flowmetry to confirm occlusion (60min) and subsequent reperfusion (**Figure 1B**). It is important to note that while the occlusion was transient, the hippocampus and other brain regions such as cortex, striatum and/or thalamus were maintained in hypoxia over the time course of the experiment (6h and 1d; **Supp. Figure 1A, B**).

**Figure 1.**
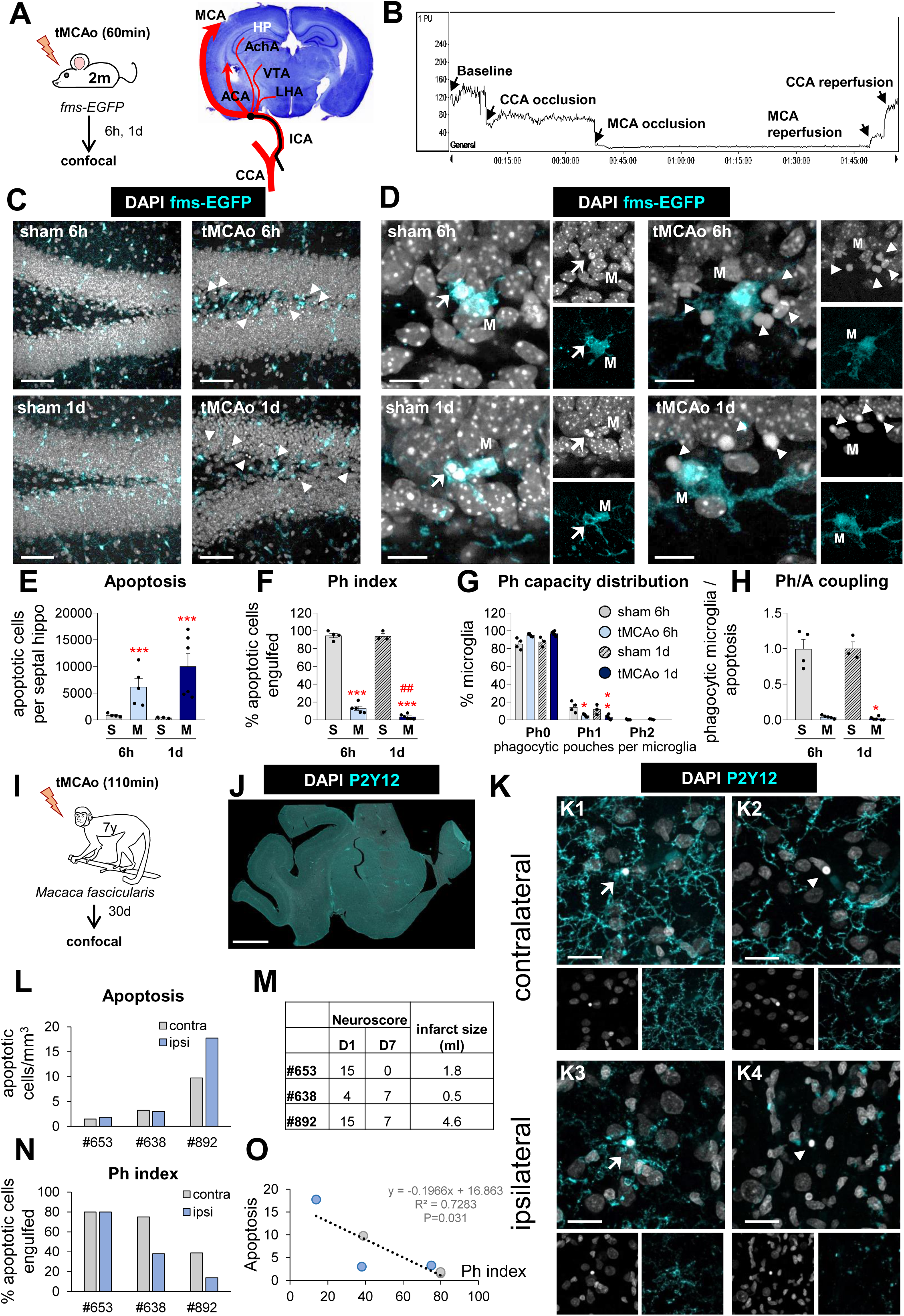
Microglial phagocytosis is impaired in mice and macaque models of tMCAo. **[A]** Experimental design of tMCAo in 2 month-old fms-EGFP mice and coronal slice with cresyl violet showing the areas irrigated by the MCA. **[B]** Laser Doppler signal graph showing CBF in the territory supplied by MCA during baseline, CCA and MCA occlusion, and reperfusion. Successful MCA occlusion, determined by CBF >70% drop from the baseline, recovers after reperfusion. The values are expressed in arbitrary Perfusion Units (PU). The area of hypoxic damage is shown in **Supp. Fig. 1A, B**. **[C]** Representative confocal z-stacks of the DG of fms-EGFP mice at 6h and 1d after tMCAo. Cell nuclei were visualized with DAPI (in white) and microglia by EGFP (in cyan). Apoptotic cells are marked with arrowheads. **[D]** Representative confocal z-stacks from the septal DG of a sham and tMCAo-treated mice at 6h and 1d, showing apoptotic cells non-phagocytosed (arrowheads) or phagocytosed (arrows) by microglia (fms-EGFP^+^, in cyan; M). **[E]** Number of apoptotic cells per septal hippocampus in sham and tMCAo. **[F]** Ph index in the septal hippocampus (% of apoptotic cells engulfed by microglia). The quantification of types of phagocytosis (ball-and-chain or apposition) are shown in **Supp. Fig. 1C**. **[G]** Histogram showing the Ph capacity of microglia (% of microglia with phagocytic pouches). The weighted Ph capacity and the number of microglia are shown in **Supp. Fig. 1D, E**. Microglial phagoptosis, radial neuroprogenitor phagocytosis, and P2Y12 expression are shown in **Supp. Fig. 2**. **[H]** Ph/A coupling (in fold change) in the septal hippocampus. Phagocytosis impairment in a model of HI is shown in **Supp. Fig. 3-5** and phagocytosis during irradiation is shown in **Supp. Fig. 6**. **[I]** Experimental design of tMCAo in *Macaca fascicularis*. Data from these animals were previously published here ^62^. Microglial phagocytosis in a tMCAo model in *Callithrix jacchus* is shown in **Supp. Fig. 7**. **[J]** Epifluorescent image of the cerebral cortex of *Macaca fascicularis* showing DAPI (nuclei, white) and P2Y12 (microglia, cyan). **[K]** Representative confocal z-stacks of the cortical regions of macaques at 30d after tMCAo from the ipsilateral (**K1, K2**) or contralateral (**K3, K4**) hemispheres. Cell nuclei were visualized with DAPI (in white) and microglia with P2Y12 (in cyan). Arrows and arrowheads point to phagocytosed (**K1, K3**) and non-phagocytosed apoptotic cells (**K2, K4**), respectively. **[L]** Number of apoptotic cells per mm^3^ in the ipsi- and contralateral hemispheres. **[M]** Table summarizing the behavioral impairment (neuroscore; higher numbers indicate more impairment) at days (D) 1 and 7 after the tMCAo, and the infarct size determined by magnetic resonance imaging (MRI) ^62^ **[N]** Ph index (% of apoptotic cells engulfed by microglia) in the ipsi- and contralateral hemispheres. **[O]** Correlation between apoptosis and Ph index in the ipsi- (light blue) and contralateral (grey) hemispheres of the three macaques. The regression line, the regression coefficient R^2^, and the adjusted p-value are shown. Bars show mean ± SEM **[E-H]**. n=3 mice (sham at 6h), n=4 mice (sham at 1d), n=5 mice (tMCAo at 6h) and n=6 mice (tMCAo at 1d). The effect of sham/tMCAo at 6h and 1d on apoptosis **[E]**, Ph index **[F]**, and microglia **[H]** was analyzed using 2-way ANOVA. Significant interactions were found between the two factors (tMCAo treatment x time); therefore, data was split into two 1-way ANOVAs to analyze statistical differences due to the time after sham/ tMCAo at each time. Holm-Sidak was used as a post hoc test. To comply with homoscedasticity, some data was Log_10_ **[E]** or (Log_10_ +1) transformed **[G, H]**. In case that homoscedasticity was not achieved with the transformation, data were analyzed using a Kruskal-Wallis ranks test, followed by Dunn method as a post hoc test **[G, H]**. (* and ^#^) represent significance compared to sham and/or tMCAo at 6h, respectively. One symbol represents p<0.05, two p<0.01, and three p<0.001. Only significant effects are shown. Scale bars=50μm, z=18.9μm **[C]**; 14μm, z=16μm **[D]**; 5mm **[J]**; 20μm **[K]**, z=18.9μm **[K1]**, 14.1μm **[K2]**, 14.1μm **[K3]**, 15.4μm **[K4]**.

We then performed immunofluorescence and confocal imaging to observe microglia, expressing the green reporter EGFP under the fms promoter (fms-EGFP mice) ^17^; and apoptotic cells, with abnormal nuclear morphology (pyknosis, karyorrhexis) using the DNA stain, DAPI (**Figure 1C, D**). Abnormal nuclear morphology is the gold standard for assessing apoptosis and, in our hands, more reliable than markers such as activated caspase 3 ^14, 15^, which has alternative functions to apoptosis. Compared to control (sham-operated mice), tMCAo mice showed more apoptotic cells (**Figure 1C**) but few cases of phagocytosis (**Figure 1D**), as determined by a microglial process forming a three-dimensional pouch that surrounded the apoptotic cell and connected to the microglial cell body ^14^. The contralateral hippocampus was not used as control because we found fluctuating levels of apoptosis. In the ipsilateral hippocampus, apoptosis increased significantly at 6h and 1d after tMCAo compared to control (**Figure 1E**), whereas the phagocytic index (Ph index, % of apoptotic cells engulfed) dropped from 94.1 ± 3.0% and 94.8 ± 2.5% in control to 13.0 ± 2.5% and 3.5 ± 1.1% in tMCAo (at 6h and 1d, respectively; **Figure 1F**). Another indication of phagocytosis impairment was that whereas in control mice phagocytosis was executed by terminal branches of microglia (“ball-and-chain” mechanism), one day after tMCAo up to 42% of phagocytosis was performed by direct apposition to the microglial soma (**Supp. Figure 1C**). We had already observed phagocytosis by apposition in models of epilepsy, where it was related to dysfunctional phagocytosis due to reduced microglial process motility and impaired apoptotic cell recognition ^15^.

To counteract the increased number of apoptotic cells, microglia would be expected to increase their phagocytic capacity (Ph capacity, % of microglia with one or more pouches) ^15^. Instead, we found that microglia reduced their Ph capacity without altering their cell numbers (**Figure 1G, Supp. Figure 1D, E**). Altogether, these changes led to a reduced coupling between phagocytosis and apoptosis (Ph/A coupling, ratio between net phagocytosis (Ph capacity x microglia) and apoptosis) after tMCAo (**Figure 1H**). In tMCAo mice, but not in control mice, we also observed occasional cases of presumptive phagoptosis, or engulfment of seemingly normal nuclei, albeit at much lower levels than apoptosis (**Supp. Figure 2A, B**). However, we did not observe compensatory phagocytosis by other presumed phagocytic cells of the hippocampus, such as astrocytes or the radial neural stem cells (rNSCs) (**Supp. Figure 2C, D**); nor by invading monocytes, identified by their lack of expression of the pan-microglial marker P2Y12 ^18^, which were not present in the hippocampus at the time points studied (**Supp. Figure 2E-G**).

We confirmed the microglial phagocytosis impairment and the lack of involvement of monocytes using a related model of hypoxia-ischemia (HI) in mice by ligation of the right common carotid artery (CCA) and exposure to 10% oxygen for 75 min ^19^. We used CX3CR1^GFP/+^/CCR2^RFP/*^ mice to discriminate microglia (RFP^-^) from monocytes (RFP^+^). As in tMCAo, microglia did not respond to the increased number of apoptotic cells with an increased Ph capacity, resulting in many apoptotic cells not phagocytosed (reduced Ph index) and disrupting the phagocytosis-apoptosis crosstalk (reduced Ph/A coupling) (**Supp. Figure 3**). In this model we observed that monocytes, which expressed CCR2 and not P2Y12 (**Supp. Figure 4**), invaded the hippocampus 3d after HI but did not engage in phagocytosis (**Supp. Figure 5A-D**). As in tMCAo, we also observed a few cases of microglial and monocyte phagoptosis (**Supp. Figure 5E, F**). The phagocytosis impairment that microglia suffered in tMCAo and HI was more evident when we analyzed the microglial response in a model of cranial irradiation ^20, 21^(**Supp. Fig. 6**). Here, mice received a single dose of 8Gy that led to a huge increase of apoptotic cells in the subgranular zone of the dentate gyrus (presumably, proliferating newborn cells, more susceptible to irradiation). In contrast to tMCAo and HI, microglia now responded by increasing phagocytosis and clearing out the dead cells within 1d. Therefore, the phagocytosis impairment observed in tMCAo and HI was not merely a saturation of the phagocytic response but rather an active phenomenon related to the pathophysiological mechanisms operating in tMCAo and HI.

We next addressed the relevance of the phagocytosis impairment for human stroke using a model of tMCAo in macaques (*Macaca fascicularis*), 30 days after the stroke ^22^ (**Figure 1I, J**). Analysis of autopsy human tissue is constrained by the postmortem delay, which affects both phagocytosis and apoptosis ^15^, microglial process motility ^23^, and microglial gene expression ^24^. We therefore analyzed the cortical temporal region of three macaques with varying levels of apoptosis (**Figure 1K, L**). Apoptosis was lower in the contralateral hemisphere compared to the ipsilateral hemisphere in the most damaged animal (#892)(**Figure 1L**), with largest infarct size and sustained behavioral deficits (**Figure 1M**). We analyzed phagocytosis using P2Y12 to label microglial processes and found that it was lower in the ipsilateral than in the contralateral hemisphere in animals #638 and #892, as determined by the Ph index (**Figure 1N**). Overall, apoptosis inversely correlated with the Ph index (**Figure 1O**), indicating a higher phagocytosis impairment in the most damaged tissue. We also analyzed cortical and hippocampal tissue from three marmoset monkeys (*Callithrix jacchus*), 45 days after tMCAo ^25^. Although we did not have proper controls, we found low absolute levels of phagocytosis, with a Ph index of 41.1% (**Supp Figure 7**). Altogether, data obtained in mice and monkey models strengthens our hypothesis that stroke impairs microglial phagocytosis.

### Energy depletion impairs engulfment and degradation via alterations in motility and lysosomal function

We then studied whether the effects of stroke on phagocytosis were related to the energy depletion induced by the lack of blood supplies, using an in vitro model of oxygen and nutrient deprivation (OND: culture medium salt solution and 1% O_2_) (**Figure 2**). In organotypic cultures, 3h and 6h of OND resulted in a phagocytosis phenotype similar to that induced by tMCAo and HI: increased apoptosis and reduced Ph index and Ph capacity, resulting in Ph/A uncoupling (**Figure 2A-E; Supp. Figure 8A-D**). Importantly, engulfment was recovered rapidly as early as 1h after reperfusion (complete medium and normoxia) indicating that the effect of energy depletion on phagocytosis was reversible in vitro. In contrast, in tMCAo we found sustained phagocytosis impairment 24h after reperfusion, possibly related to the buildup of hypoxia over time (**Supp. Figure 1A, B**), which has been associated with the death in rigor of pericytes and the irreversible constriction of brain capillaries ^26^. The reduced engulfment induced by OND was likely related to a reduction in microglial process motility, as determined by 2-photon microscopy in organotypic slices under OND (**Figure 2F-I**).

**Figure 2.**
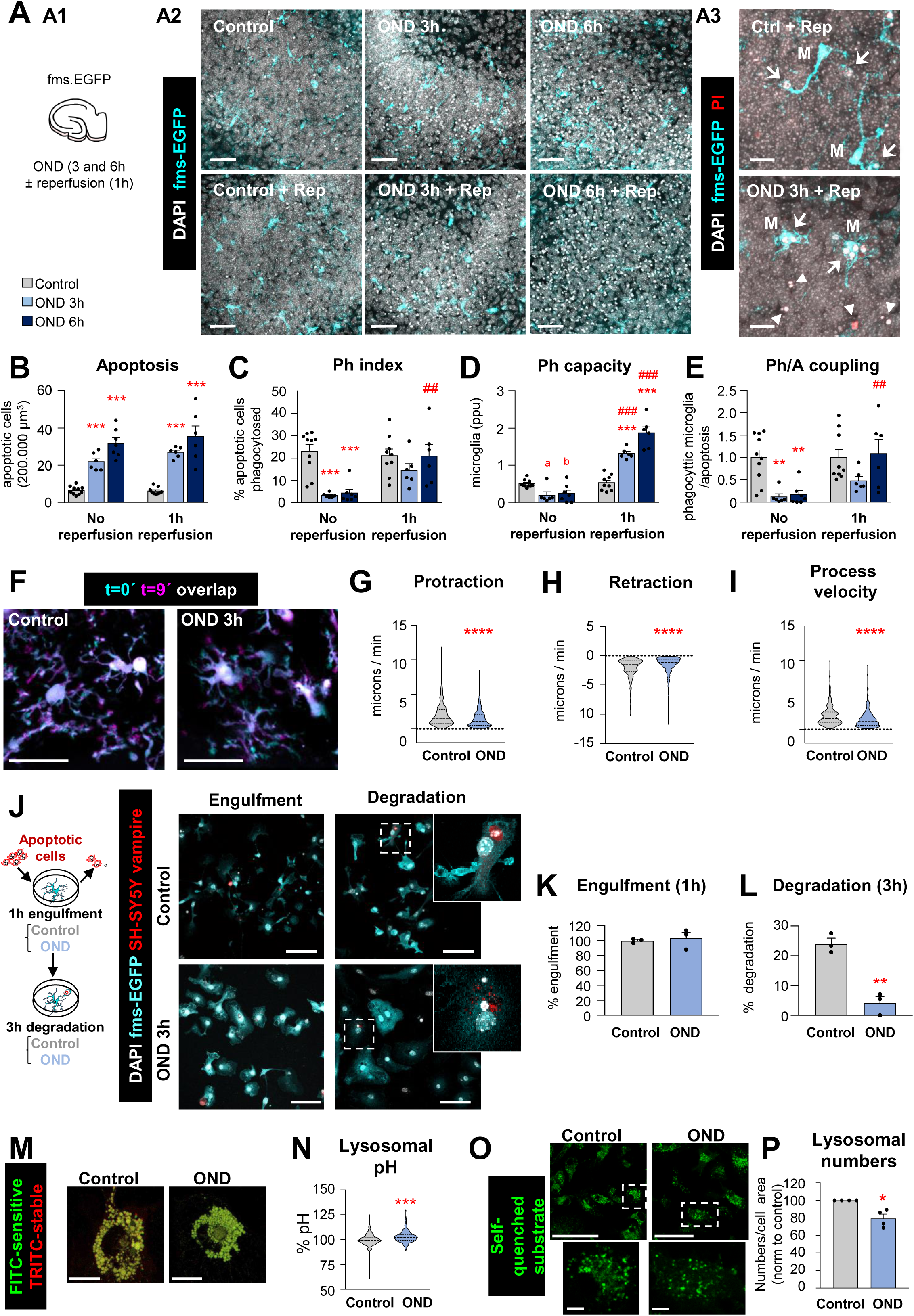
Engulfment and degradation of apoptotic cells are impaired after OND due to alterations in motility and lysosomal function. **[A]** Experimental design showing the exposure of hippocampal organotypic slices (fms-EGFP) to OND (3 and 6 hours) in the presence and absence of 1h reperfusion **[A1]**. Representative confocal images of the DG after OND **[A2]**. Normal or apoptotic (pyknotic/karyorrhectic) nuclear morphology was visualized with DAPI (white) and microglia by the transgenic expression of fms-EGFP (cyan). Under reperfusion conditions, membrane permeability (characteristic of necrotic cells) was observed with PI (red). High magnification images show primary apoptotic cells (pyknotic/karyorrhectic, PI^-^) or secondary necrotic cells (pyknotic/karyorrhectic, PI^+^) engulfed (arrows) or not-engulfed (arrowheads) by microglia (M) (EGFP^+^) **[A3]**. Secondary necrotic cells were very few and were pooled together with the rest of pyknotic/karyorrhectic cells and labeled as apoptotic thereafter. Single channel images of **A3** are shown in **Supp. Figure 8A**. **[B]** Number of apoptotic cells in 200.000µm^3^ of the DG. **[C]** Ph index (% of apoptotic cells phagocytosed by microglia). **[D]** Weighted Ph capacity (number of phagocytic pouches containing an apoptotic cell per microglia, in parts per unit (ppu). The Ph capacity histogram and the number of microglia is shown in **Supp. Figure 8B-D**. **[E]** Ph/A coupling expressed as fold-change; ratio between net phagocytosis and total levels of apoptosis. **[F]** Representative projections of 2-photon images of microglial cells at t0 (cyan) and t9 (magenta) from hippocampal organotypic slices (CX3CR1^GFP/+^) under control and OND conditions. **[G, H, I]** Microglial process motility: protraction **[G]**, retraction **[H]**, and process velocity **[I]**. **[J]** Experimental design of the phagocytosis assay to assess engulfment and degradation of apoptotic cells under control and OND conditions. A table summarizing the treatments is shown in **Supp. Fig. 8E.** Representative images of primary microglia fed with apoptotic SH-SY5Y vampire cells during engulfment and degradation. Nuclei were visualized with DAPI (white), microglia by expression of EGFP (cyan), and SH-SY5Y neurons by expression of the red fluorescent protein Vampire. **[K, L]** Percentage of phagocytic microglia after engulfment (1h) and degradation (3h after engulfment). Only particles fully enclosed by microglia were identified as being phagocytosed. Raw % of phagocytic microglia are shown in **Supp. Fig. 8F**. Phagocytosis under more stringent conditions (shorter incubation, fewer apoptotic cells) is shown in **Supp. Fig. 8G**. **[M]** Representative confocal images of microglia incubated with dextran molecules conjugated to two fluorophores: FITC (pH sensitive) and TRITC (pH stable) located in the lysosomes, whose ratio serves as an indirect measurement of the lysosomal pH. **[N]** Lysosomal pH expressed as % normalized to control values. Note the truncated Y axis. **[O]** Representative confocal images of microglia loaded with the self-quenched substrate. **[P]** Percentage of lysosomal numbers normalized to control values under control and OND conditions. The cytoplasmic area occupied by lysosomes and the lysosomal activity is shown in **Supp. Figure 8H, I**. Bars show mean ± SEM **[B-E, K, L]**. Violin plots show the data distribution including extreme values; lower and upper hinges correspond to the first and third quartile respectively **[G, H, I, N]**. n = 6-10 mice per group **[B-E]**; n=355 processes from 98 cells from 12 animals (control), and n=222 processes from 57 cells from 9 animals (OND) [G-I]; n=3 independent experiments **[K, L]**; n=530 cells (control) and 452 cells (OND) from 4 independent experiments **[N]**. n=4 independent experiments **[P]**. Data was analyzed by two-way ANOVA followed by Holm-Sidak post hoc tests **[B]**. When an interaction between factors was found, one-way ANOVA (factor: treatment) was performed followed by Holm-Sidak post hoc tests **[C-E]**. To comply with homoscedasticity, some data was Log_10_ transformed **[B]** or Ln transformed **[E]**. Other data was analyzed by Kruskal-Wallis rank test **[G-I]** or by Student’s t-test. **[J, K, L]**. (*and ^#^) represent significance between control and OND, or no reperfusion vs reperfusion, respectively: one symbol represents p<0.05, two symbols represent p<0.01, three symbols represent p<0.001, and four symbols represent p<0.0001; (a) represents p=0.06 and (b) represents p=0.07 (control vs OND). Scale bars=50µm, z=10.5µm **[A2]**;15 µm, z=16.8µm **[A3]**, 20µm, z=22μm [**F**]; 5µm, z=8.5μm **[J]**, 7µm **[M]**; 50µm low magnification, 5µm high magnification, z= single plane **[O]**.

To identify the mechanisms of phagocytosis impairment during energetic depletion we used primary cultures of microglia co-incubated with a neuronal cell line in which we induced apoptosis (SH-SY5Y-Vampire pre-treated with staurosporine). We analyzed the amount of microglia with Vampire^+^ particles at two time points to discriminate engulfment (1h of co-incubation) and degradation (3h after washout) in cells treated with OND (**Figure 2J; Supp. Figure 8E**). This model uncovered an effect of OND on degradation (**Figure 2K, L, Supp. Figure 8F**), which could not be assessed in vivo or in organotypic cultures because it is downstream of engulfment. Nonetheless, our in vitro model did not reveal the expected effect on engulfment, not even when we co-incubated microglia in more stringent conditions (fewer apoptotic SH-SY5Y cells, or reduced co-incubation time)(**Supp. Figure 8G**), possibly because in vitro models of phagocytosis do not fully mimic the complexity of the “find-me” and “eat-me” signals that regulate engulfment in vivo ^27^. We therefore focused on understanding the basis of degradation impairment by studying lysosomes, the degradative organelles. We found that under OND microglia had a small but significant increase in lysosomal pH **(Figure 2M, N)** and reduced lysosomal number (**Figure 2O, P**). Although individual lysosomes had a similar enzymatic activity (**Supp. Figure 8H, I**), these changes resulted in a reduced degradative capacity of microglia under OND. In conclusion, the energetic depletion associated with stroke activated several cell processes that affected both engulfment and degradation of apoptotic cells that included alterations in motility and in lysosomal pH and number.

### Autophagy induction after OND

The lysosomal alterations induced by OND motivated us to focus on autophagy, a major intracellular degradative pathway with several reported alterations in models of stroke ^28^. Autophagy and phagocytosis are relatively similar processes whose goal is to degrade intracellular (autophagy) or extracellular (phagocytosis) cargo, and they share some mechanisms and cellular machinery ^29^. To study the autophagy status in microglia we used three complementary approaches (**Figure 3**): analysis of the autophagosome marker LC3 by confocal imaging and western blot, and direct visualization of autophagosomes by transmission electron microscopy (TEM). First, we used a tandem plasmid GFP-RFP-LC3 to transfect the microglial cell line BV2 (**Figure 3A-C**). Cytoplasmic LC3 was observed as a diffuse signal, whereas autophagosome LC3 was located in puncta. As GFP fluorescence is pH-sensitive (whereas RFP is pH-stable), a reduced GFP/RFP fluorescence ratio indicated the fusion of the autophagosomes with lysosomes (autolysosomes) ^30^. Microglial cells under OND, similar to cells treated with the autophagy inducer rapamycin, showed a trend to increased RFP intensity in puncta (**Figure 3B, C1**), suggesting increased LC3 presence in autophagic vesicles. However, GFP intensity did not increase proportionally **(Figure 3B, C2)** and the GFP/RFP ratio was reduced in both OND and rapamycin-treated cells compared to the control group **(Figure 3B)**, suggesting increased LC3 degradation in autolysosomes. Similar to rapamycin treatment, OND tended to decrease both the number and area of LC3 puncta **(Figure 3C3, C4)**, although significant changes were not evident due to the high variability of data. Overall, these results indicate that OND and rapamycin enhance autophagy flux in microglia.

**Figure 3.**
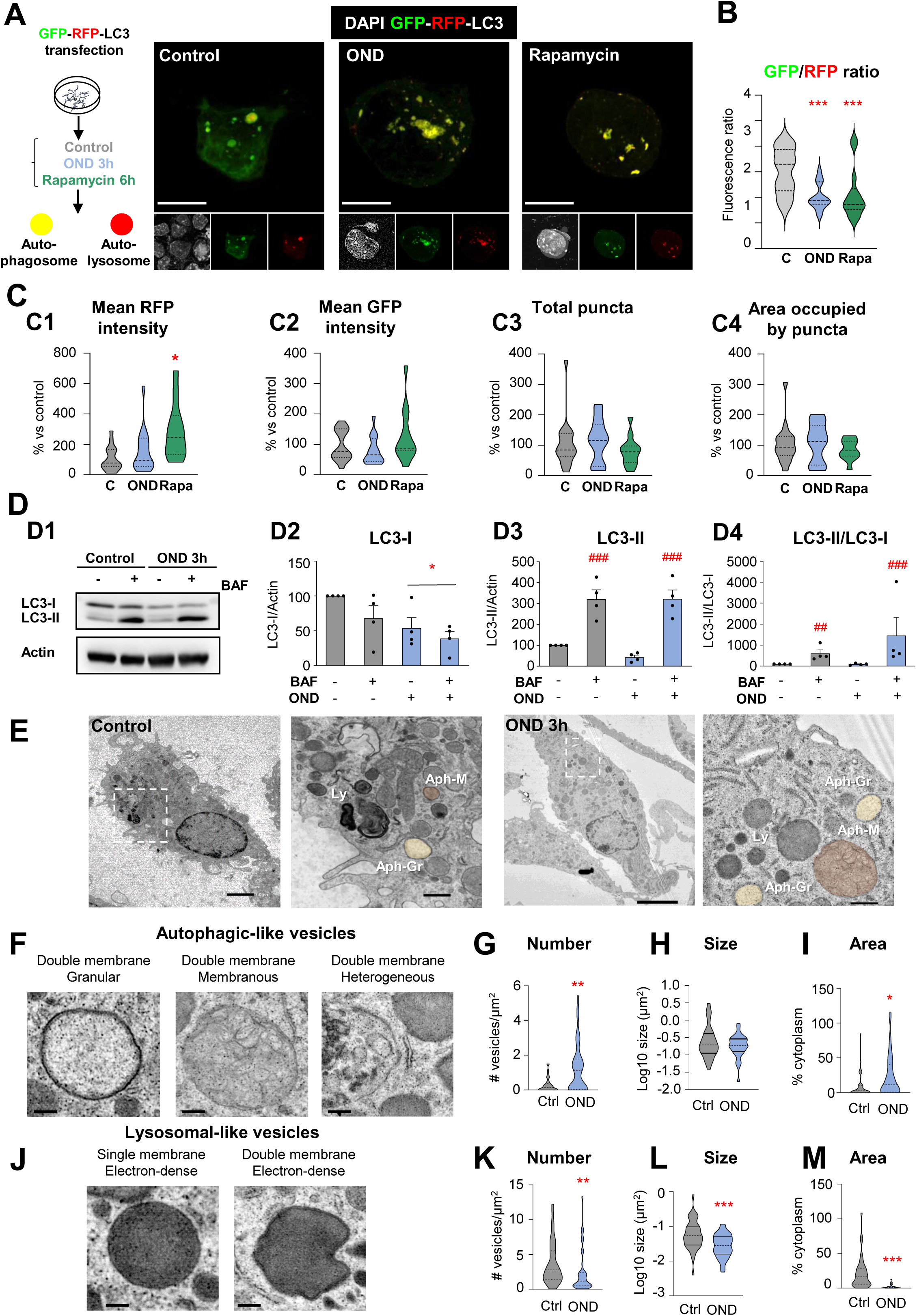
The autophagy compartment is remodeled after OND. **[A]** Experimental design used to transfect BV2 microglia-like cells with the fluorescent tandem GFP-RFP-LC3 to assess autophagy flux in control conditions (C) and after OND (3h) or rapamycin (Rapa, 100nM, 6h) treatments. Representative confocal images of control, OND and rapamycin treated microglia. Nuclei are stained with DAPI (white), autophagosomes and autolysosomes are differentiated according to the tandem expression (yellow and red, respectively). **[B]** GFP/RFP mean fluorescence intensity ratio, indicative of autophagy flux. **[C]** Mean RFP fluorescence intensity **[C1],** mean GFP fluorescence intensity **[C2],** total number of puncta **[C3]**, and area occupied by puncta **[C4]** per cell and normalized to control conditions (expressed as % change versus control conditions). **[D]** Primary microglia were exposed to OND for 3 hours in the presence and absence of bafilomycin-A1 (BAF, 100 nM, 3 hours) to assess autophagy flux by LC3 turnover assay. Delipidated (∼1 KDa) and lipidated (∼17 KDa) LC3 levels were analyzed by western blot. Beta-actin (∼42 KDa) was used as a loading control. Representative blots showing LC3-I, LC3-II and actin bands **[D1]**, LC3-I levels normalized to actin **[D2]**, LC3-II levels normalized to actin **[D3]**, LC3-II levels normalized to LC3-I levels **[D4]**. BV2 autophagy flux is shown in **Supp. Figure 8J**. **[E]** Representative transmission electron microscopy images of primary microglia in control and OND conditions. Ly: lysosomes; Aph-Gr: autophagosomes with granular cargo (yellow); Aph-M: autophagosomes with membranous cargo (orange). **[F]** Details of autophagic-like vesicles identified as containing at least a portion of double membrane with different types of cargo (granular, membranous, heterogeneous). **[G-I]** Quantification of autophagic-like vesicle number per µm^2^ **[G]**, size in µm^2^ (in logarithmic scale) **[H]**, and percentage of cytoplasm occupied **[I]**. **[J]** Details of lysosomal-like vesicles identified as electron-dense vesicles with single or double membrane. **[K-M]** Quantification of lysosomal-like vesicle number per µm^2^ **[K]**, size in µm^2^ (in logarithmic scale) **[L]**, and percentage of cytoplasm occupied **[M]**. Bars show mean ± SEM **[D]**. Violin plots show the data distribution, including extreme values; lower and upper hinges correspond to the first and third quartile, respectively **[B-C, G-I, K-M]**. n=12-17 cells from 3 independent experiments **[B-C]**; n=4 independent experiments **[D]**; n=36-38 cells from 3 independent experiments **[G-I, K-M]**. Data was analyzed by one-way ANOVA followed by Bonferroni post hoc test **[B]**, by Kruskal-Wallis one way ANOVA on ranks followed by Dunn’s multiple comparisons **[C]** and by two-way ANOVA followed by Holm-Sidak post hoc test **[D].** Other data was analyzed by non-parametric Mann-Whitney test **[G-I, K-M]**. (* and ^#^) represent significance between control and OND or rapamycin, or between bafilomycin and no bafilomycin, respectively: one symbol represents p<0.05, two symbols represent p<0.01 and three symbols represent p<0.001. Scale bars=10µm, z=1.9µm (control), 3.3µm (OND), and 3.9µm (Rapamycin) **[A]**; 2µm (control), 5µm (OND), 500nm (high magnification) **[E]**; 500nm **[F, J]**.

This effect was, however, not evident by western blot of LC3, which differentiates LC3-I (unconjugated, cytosolic) from LC3-II (conjugated to autophagosomal lipids). Primary microglia (**Figure 3D**) or BV2 cells (**Supp. Figure 8J**) were maintained in OND in the presence or absence of the lysosomal inhibitor bafilomycin to assess the autophagic flux ie., the accumulation of autophagosomes ^31^. We detected a reduction in LC3-I that was more evident in BV2 cells (**Figure 3D2, Supp. Figure 8J2**) suggestive of LC3-I lipidation into autophagosomes. Indeed, LC3-II/LC3-I tended to increase after OND (**Figure 3D4, Supp. Figure 8J4**), reaching significance in BV2 cells (**Supp. Figure 8J4**). However, no significant effects were observed on LC3-II/actin accumulation after addition of bafilomycin as would be expected in a classic autophagy induction (**Figure 3D3, Supp. Figure 8J3).** We attribute these inconclusive results to a lower sensitivity of western blot compared to imaging techniques in detecting autophagy induction, at least in our hands. Finally, we used TEM to directly visualize autophagosomes (double membrane vesicles with granular, membranous or heterogeneous contents) and lysosomes (electrondense vesicles) in primary microglia (**Figure 3E-M**). Control microglia had evident autophagosomes, suggestive of ongoing autophagy in basal conditions (**Figure 3E**). After OND, we found more autophagosomes and they occupied a larger area of the cytoplasm (**Figure 3F-I**), supporting the induction of autophagy after OND. In addition, we found a reduced number of smaller lysosomes that occupied less cytoplasm (**Figure 3J-M**), in agreement with the increased pH and reduced lysosomal number (**Figure 2M-P**). We speculated that their fusion with autophagosomes would lead to their consumption in the short-term, as has been observed before ^32^, resulting in reduced numbers and a reduced degradative capacity available for phagocytosis. This idea prompted us to study the functional relationship between autophagy and phagocytosis in basal conditions and during energy depletion.

### Basal autophagy sustains microglial physiology, including phagocytosis

We first focused on basal autophagy and used transgenic mice deficient in several autophagic genes to assess their impact on phagocytosis (**Figure 4**, **Supp. Figure 9A**): the LC3 protease ATG4B; and the phagophore extending class III PI3K proteins Beclin-1 and AMBRA1. In the hippocampus of mice deficient in ATG4B, we found a reduction of phagocytosis (Ph index) and the number of microglia (**Figure 4A-E**). Similarly, in mice with inducible Beclin1 deficiency in microglia (driven by TMEM119), we found a tendency to reduced phagocytosis (**Supp. Figure 9B-E**). In contrast, we found no effect on phagocytosis in mice with heterozygous deficiency in AMBRA1 (**Supp. Figure 9F-I**). We did not use AMBRA1 full KO mice because they are embryonically lethal ^33^. This data suggests that basal autophagy, which may be controlled by a specific set of genes in microglia compared to other cell types ^34^, is necessary for an efficient phagocytosis.

**Figure 4.**
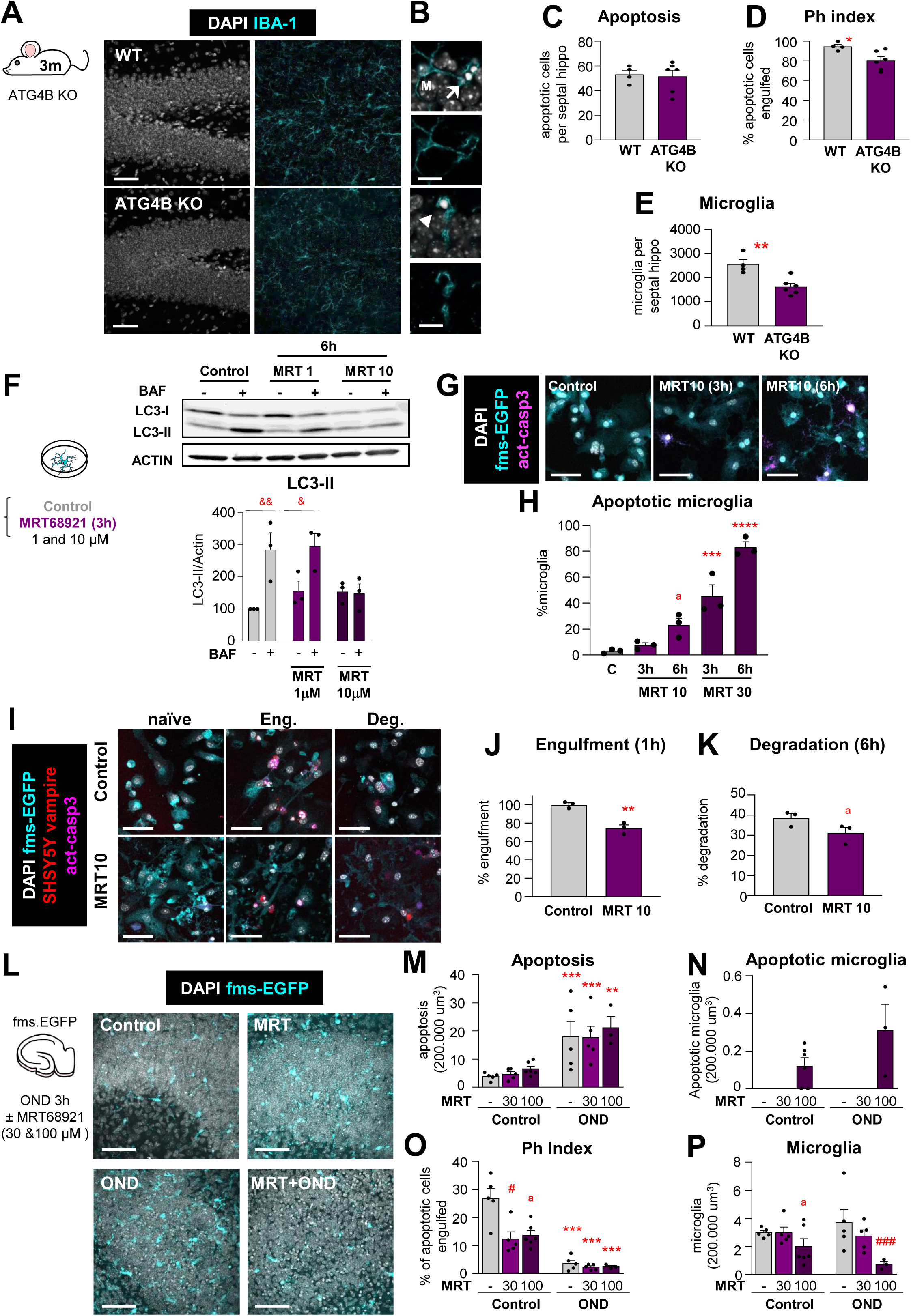
Basal autophagy is essential for microglial survival and function. **[A]** Representative confocal z-stacks of the DG of 3 month-old wild-type (WT) and ATG4B knock-out (ATG4B KO) mice. Healthy or apoptotic nuclei (pyknotic/karyorrhectic) were visualized with DAPI (white) and microglia were stained for IBA-1 (cyan). A cartoon representing the proteins involved in the autophagic response is shown in **Supp. Figure 9A**. Phagocytosis in TMEM-Beclin KO and AMBRA^+/-^ mice are shown in in **Supp. Figure 9B-E** and **F-I**, respectively. **[B]** High magnification examples of phagocytosed (arrows) and nonphagocytosed (arrowheads) apoptotic cells. **[C]** Number of apoptotic cells per septal hippocampus in WT and ATG4B KO mice. **[D]** Ph index in the septal hippocampus (% of apoptotic cells engulfed by microglia). **[E]** Number of microglial cells per septal hippocampus in WT and ATG4B KO mice. **[F]** Experimental design of the dose-response administration of Ulk1/2 inhibitor MRT68921 to primary microglia. Representative blot showing relative levels of LC3-I and LC3-II after 1 and 10μM MRT68921 administration for 6h. Quantification of the LC3-II levels (referred to actin) after 1 and 10μM MRT68921 in the presence and absence of the lysosomal inhibitor, bafilomycin A (BAF, 100 nM). The quantification of LC3-I is shown in **Supp. Fig. 10A, B**. The analysis of LC3-I and II after 3h of MRT68921 (10 and 30μm) is shown in **Supp. Fig. 10C**. This data is reprinted with permission from Frontiers in Immunology ^31^. **[G]** Representative confocal images of primary fms-EGFP microglia treated with MRT68921 (10 and 30μM). Nuclei were visualized with DAPI (white), microglia with their constitutive EGFP expression (cyan), and apoptotic cells with activated caspase 3 (act-casp3^+^, magenta). Images of all experimental groups are shown in **Supp. Fig. 10D**. **[H]** Percentage of apoptotic microglia assessed by their healthy or apoptotic nuclei (pyknotic/karyorrhectic). **[I]** Representative confocal images of naïve (non-phagocytic), engulfing and degrading fms-EGFP microglia (cyan), after the addition of apoptotic SH-SY5Y vampire with RFP (red); nuclear morphology (pyknotic/karyorrhectic) was assessed with DAPI (white). A table summarizing the treatments is shown in **Supp. Fig. 11A**. **[J, K]** Percentage of phagocytic microglia after engulfment (1h) and degradation (6h after engulfment). Only particles fully enclosed by microglia were identified as phagocytosis. Raw % of phagocytic microglia is shown in **Supp. Fig. 11B** and the % of apoptotic microglia is shown in **Supp. Fig. 11C**. **[L]** Experimental design and representative confocal images of the DG after treatment with MRT68921 (100 µM) for 3 hours in the presence and absence of OND. Normal or apoptotic (pyknotic/karyorrhectic) nuclear morphology was visualized with DAPI (white) and microglia by the transgenic expression of fms-EGFP (cyan). **[M]** Number of apoptotic cells in 200.000 µm^3^ of the DG. **[N]** Number of apoptotic microglia. Apoptotic microglia were discriminated from apoptotic cells contained in microglial pouches thanks to their expression of EGFP within the nuclei and the lack of a process connecting it to a healthy microglial soma. An example is shown in **Supp. Fig. 11D**. **[O]** Ph index (% of apoptotic cells phagocytosed by microglia). **[P]** Number of microglia in 200.000 µm^3^ of the DG. The weighted Ph capacity and distribution, and the Ph/A coupling are shown in **Supp. Fig. 11E-G**. Bars show mean ± SEM. n=4-6 mice per group [**C-E**], n=3 independent experiments [**F, H, J, K**], n=3-6 mice per group **[M, N, O, P]**. Data was analyzed using Student’s t-test analysis **[C-E, J-K],** by two-way ANOVA followed by Holm-Sidak post hoc tests **[F]** after logarithmic transformation **[ M, O, P]**, or by 1-way ANOVA followed by Tukeýs multiple comparisons **[H]**. (&) represents significance between bafilomycin-treated and non-treated groups: one symbol represents p<0.05 and two symbols represent p<0.01. (* and ^#^) represent significance compared to the control group and between MRT-treated vs MRT-untreated, respectively: one symbol represents p<0.05, two symbols represent p<0.01, three symbols represent p<0.001 and four symbols represent p<0.0001; (a) represents significance between MRT-treated vs MRT-untreated groups in control conditions; p=0.051 **[H],** p=0.1080 **[K]**, p=0.055 **[O]** and p=0.127 **[P]**. Scale bars= 50μm, z=36.4μm [**A**]; 10μm **[B]**; 50μm, z=8.5μm [**G, I**]; 50 µm and z=11.2µm **[L]**.

We further confirmed the functional relationship between autophagy and phagocytosis with MRT68921, an inhibitor of the upstream autophagy regulator ULK1/2 (**Figure 4F-K; Supp. Figure 9A**). Here, western blot of LC3 was sufficiently sensitive to detect a blockade of autophagy flux in primary cultures (**Figure 4F, Supp. Figure 10A-C**). While high concentrations of MRT led to microglial apoptosis (**Figure 4G, H**), we identified a dose of MRT (10μM, 6h) that blocked autophagy without inducing microglial apoptosis (**Figure 4F-H, Supp. Figure 10A-D**) and used it to test its effects on phagocytosis (**Supp. Figure 11A**). MRT reduced very effectively apoptotic cell engulfment and showed a trend to reduced degradation (**Figure 4I-K; Supp. Figure 11A-C**). Therefore, autophagy inhibition with MRT impaired both survival and phagocytosis in vitro, confirming the effects of ATG4B deficiency in vivo.

Additionally, we used organotypic cultures to test the effect of autophagy inhibition with MRT on both basal and OND-treated cultures (**Figure 4L-P**). We used a higher concentration of MRT (30μM and 100μM) to account for the drug diffusivity throughout the tissue slice. At these concentrations, MRT did not induce global apoptosis (**Figure 4L, M**). In control conditions, MRT induced a small amount of microglial apoptosis (**Figure 4N**) and reduced phagocytosis (**Figure 4O; Supp. Figure 11E-G**), further supporting the relevance of basal autophagy on microglial health and phagocytosis efficiency. Under OND, MRT led to a stronger induction of microglial apoptosis (**Figure 4N; Supp. Figure 11D**) and to reduced microglial numbers (**Figure 4P**), indicating that the autophagy induction after OND was protective for microglia. However, MRT did not further enhance the OND-induced phagocytosis impairment (**Figure 4O; Supp. Figure 11E-G**), showing that either inhibiting autophagy or OND led to a similar phagocytosis blockade.

### Rapamycin prevents phagocytosis impairment

Finally, we tested the opposite strategy and induced autophagy using rapamycin (**Figure 3A-C**) to address whether it could prevent phagocytosis dysfunction during tMCAo in vivo and OND in vitro (**Figure 5**). We selected rapamycin because of its beneficial reported effects in several brain disease models ^28^, including models of stroke ^35^, and because while it is largely known as an inhibitor of the mTORC1 pathway ^36^, it also promotes lysosomal biogenesis ^37, 38^ thus enhancing autophagy at different levels of the process (**Supp. Figure 9A**). Mice were pretreated for two consecutive days with rapamycin (10mg/kg) ^38^ or vehicle before being exposed to tMCAo, and were sacrificed 6h later (**Figure 5A**). To control for surgery heterogeneity, in each surgery session a pair of one vehicle and one rapamycin mouse were operated. All mice had MCA occlusion and reperfusion confirmed with laser Doppler flowmetry (**Figure 5B**). At the time point tested, rapamycin had no overall effect on the tMCAo-induced damage, as it had no effect on hippocampal apoptosis or on microglial numbers (**Figure 5C-E**). Importantly, however, rapamycin partially recovered phagocytosis globally (**Figure 5F**) and in 5/6 mouse pairs (**Figure 5G**), although the Ph index did not reach the expected values of untreated animals (**Figure 1F**).

**Figure 5.**
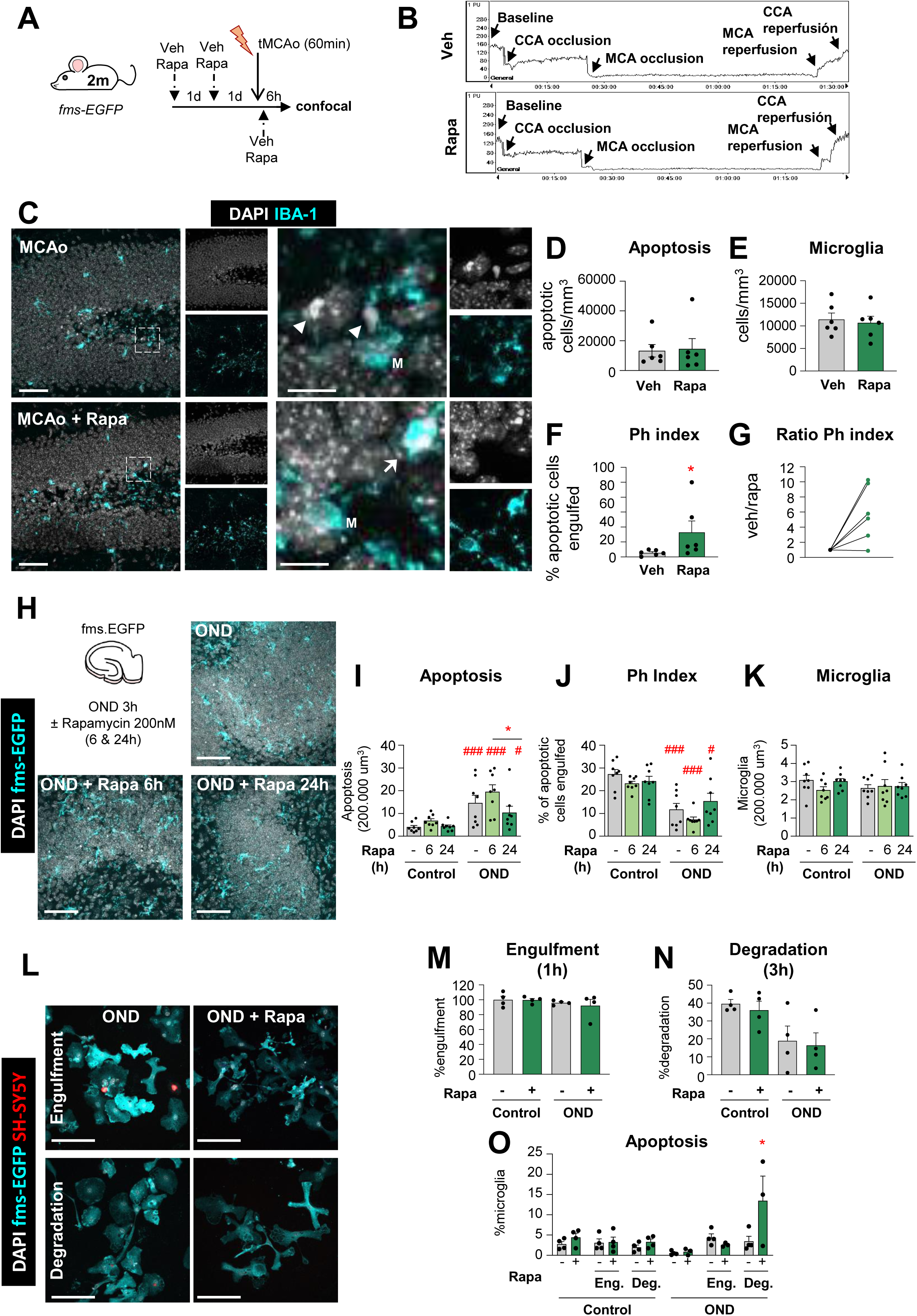
Rapamycin reverts the tMCAo-induced phagocytic dysfunction in vivo but not in vitro. **[A]** Experimental design showing the daily administration of rapamycin (10mg/kg, ip) two days prior to the tMCAo in 2 month-old fms-EGFP mice. Mice received a third rapamycin injection right after reperfusion, and were sacrificed 6h later. **[B]** Representative laser Doppler signal graph showing the effective MCA occlusion and reperfusion in both vehicle and rapamycin-treated mice. **[C]** Representative confocal z-stacks of the DG of fms-EGFP mice 6h after tMCAo, treated with vehicle or rapamycin (10mg/kg, ip). Cell nuclei were visualized with DAPI (in white) and microglia (fms-EGFP^+^, in cyan). Arrowheads point to nonphagocytosed apoptotic cells and arrows to phagocytosed apoptotic cells. M labels a microglial soma. **[D]** Density of apoptotic cells in the septal hippocampus. **[E]** Density of microglial cells in the septal hippocampus. **[F]** Ph index in the septal hippocampus (% of apoptotic cells engulfed by microglia). **[G]** Normalized ratio of Ph index change in each rapamycin-treated mice over its same day vehicle-treated mice. **[H]** Experimental design and representative confocal images of hippocampal organotypic cultures treated with vehicle or rapamycin (200nM; 6 and 24h) exposed to OND (3h). Normal or apoptotic (pyknotic/karyorrhectic) nuclear morphology was visualized with DAPI (white) and microglia by the transgenic expression of fms-EGFP (cyan). Rapamycin- and vehicle- treated control cultures are shown in **Supp. Fig. 12A**. **[I]** Number of apoptotic cells in 200.000 µm^3^ of the DG. **[J]** Ph index (% of apoptotic cells phagocytosed by microglia). The weighted Ph capacity and the Ph/A coupling are shown in **Supp. Fig. 12B, C**. **[K]** Number of microglia in 200.000 µm^3^ of the DG. [**L**] Representative confocal images of microglia (fms-EGFP^+^, cyan) engulfing (1h) and degrading (3h) apoptotic SH-SY5Y vampire neurons (red) under OND conditions in the presence of rapamycin. Nuclear morphology was assessed with DAPI (white). Only particles fully enclosed by microglia were identified as phagocytosis. A table summarizing the treatments is shown in **Supp. Fig. 12D**, the effect of rapamycin in LC3 levels is shown in **Supp. Fig. 12E,** and control cells are shown in **Supp.Fig. 12F**. [**M, N**] Percentage of phagocytic microglia after engulfment (1h) and degradation (3h after engulfment). [**O]** Percentage of apoptotic (pyknotic/karyorrhectic) microglia after the phagocytosis assay and rapamycin treatment in both control and OND. Bars show mean ± SEM. Dot and line plot represents the normalized ratio between rapamycin- and vehicle- treated animals (by pairs). n=6 mice per group **[D-G]**, n=8 mice per group **[I-K]**, and n=4 independent experiments **[M, N]**. Data was analyzed using a Student’s t-test **[D-G]**; by two-way ANOVA followed by Holm-Sidak post hoc tests when appropriate **[I-K, M-N]**; or by one-way ANOVA followed by Tukeýs post hoc test when a significant interaction in two-way ANOVA was found **[O]**. Some data (**[I]**) was Log_10_ transformed to comply with homoscedasticity. Asterisks represent significance between untreated and rapamycin-treated mice or cultures: (*) represents p<0.05. (#) represent significance between OND and control cultures: # represents p<0.05, and ### represents p<0.001. Scale bars=50µm, z=19.6µm; inserts bar=10µm z=9.8µm **[C]**; 50µm, z=11.2µm **[H]**; 50µm, z=8.5µm **[L]**.

This partially protective effect on phagocytosis was however not recapitulated in organotypic cultures (**Figure 5H-K; Supp. Figure 12A-C**). Here, rapamycin reduced global apoptosis in the longest time point tested (24h, 200nM) but had variable effects on phagocytosis that were not significant. Similarly, rapamycin had no effect on either engulfment or degradation in primary cultures (**Figure 5L-N; Supp. Figure 12D**) at a concentration that induced autophagy (100nM), as observed by tandem GFP-RFP-LC3 in BV2 cells (**Figure 3A-B)** but not by LC3 western blot in primary microglia (**Supp. Figure 12E**). In addition, we detected a detrimental effect of rapamycin in phagocytic (degradative) microglia in OND conditions that lead to their demise (**Figure 5O; Supp. Figure 12F**), possibly related to an over-stimulation of microglial autophagy in vitro. While we cannot disregard the possibility that the time points and concentrations were not optimized in vitro or that different pathways regulate microglial autophagy in vitro and in vivo, another possibility to explain the discrepant effects of rapamycin after tMCAo and OND is that the partial protection afforded by rapamycin in tMCAo mice was not due to a direct effect on microglia. Thus, while blocking autophagy was clearly detrimental, enhancing autophagy in microglia proved to be more challenging. Overall, our results confirm that the phagocytosis impairment can be modulated in vivo and is a druggable target to be explored.

## DISCUSSION

In this paper, we demonstrate that microglial phagocytosis is an early therapeutic target in stroke, amenable for drug interventions. We first show evidence of phagocytosis dysfunction in mouse and monkey models of stroke, both in vivo and in vitro. Next, we reveal some of the underlying cellular mechanisms that affect both engulfment and degradation of apoptotic cells, including reduced microglial process motility and lysosomal alterations. Then we show that the energetic stress associated with stroke leads to increased autophagy, whose basal levels are essential for microglial survival and function both in vivo and in vitro. Finally, we demonstrate that the autophagy inducer rapamycin partially prevents the stroke-induced phagocytosis impairment in vivo. Overall, we shed light onto two unappreciated microglial activities with key roles during stroke and high therapeutic potential: autophagy, responsible for intracellular recycling, and the celĺs well-being and function; and phagocytosis, responsible for extracellular laundering and controlling inflammatory responses.

### Exploiting microglial phagocytosis as a future therapeutic target

The profits of microglial phagocytosis for the diseased brain are evident: prevention of intracellular content spillover and immunomodulatory effects ^10^. However, its therapeutic potential has been largely unappreciated, likely because phagocytosis was presumed to occur rather than directly assessed ^27^. Here we have used a quantitative approach that has allowed us to discover microglial phagocytosis dysfunction in stroke, similar to what we had observed in mouse and human epilepsy ^15, 16^. In these diseases, it is necessary not only to prevent the death of neurons, but also to accelerate the removal of neuronal debris by developing new strategies to harness phagocytosis.

Pioneer work in cancer has catapulted macrophage phagocytosis as a consolidated target with several ongoing clinical trials ^39^. Here, deficient phagocytosis is due to tumor cells escaping recognition by macrophages. As such, most efforts have been put into developing opsonizing antibodies to coat the tumor cells and facilitate their interaction with macrophages ^40^. Another rising idea is to interfere with phagocytosis checkpoints such as CD47, a “dońt-eat-me” signal that interacts with SIRPα receptors on the macrophage ^41^. In the brain, however, CD47 expressed by healthy synapses prevents their excessive phagocytosis by microglia, at least during development ^42^. Unhinged phagocytosis could lead to phagoptosis, i.e., the engulfment of stressed but viable cells ^43^, responsible for delayed neuronal death in stroke models ^44^. These examples highlight that translating cancer-based approaches into effective brain therapies may not be straightforward and requires a deeper understanding of the mechanisms operating on microglial phagocytosis dysfunction, as we will discuss next.

### Impairment of microglial phagocytosis during stroke

One key difference between cancer and the diseased brain is that in the first case, tumor cells are the ones to blame, as they develop mechanisms to escape phagocytosis. In contrast, in stroke the problem does not lie on the target cells but on the phagocytes, as microglial function is severely compromised due to the energy depletion. We found that the lack of oxygen and nutrients reduced the motility of microglial processes and interfered with the engulfment phase of phagocytosis. It also altered the microglial lysosomal number, pH, and the celĺs degrading capacity, which was likely related to reduced apoptotic cell degradation. Finally, it induced autophagy, a cell process that we found necessary for microglial survival and phagocytosis. In addition, environmental-related factors are also likely to play a role on microglial phagocytosis dysfunction. An example is extracellular ATP, one of the major “find-me” signals from apoptotic cells sensed by purinergic receptors on microglia ^45^. However, ATP is also a neurotransmitter, widely released during pathological conditions such as epilepsy and ischemia ^46^. These two sources of ATP put microglia in conflict and disrupt their targeting of apoptotic cells, resulting in impaired phagocytosis during epilepsy ^15^ and possibly during stroke. Inhibitors of the purinergic receptor P2Y12, such as clopidogrel are currently used as anticoagulants in several cardiovascular diseases ^47^, but a side effect not considered is their inhibitory action on microglial phagocytosis ^27^. In sum, this trio of mechanisms that relate to the target, the phagocyte, and/or the environment should be considered when designing effective therapies to recover or potentiate phagocytosis.

Here, we have used rapamycin to prove the principle that phagocytosis dysfunction during stroke can be modulated in vivo. The autophagy inducer rapamycin has become an increasingly popular drug since its discovery in the early nineties in soil samples from Easter Island ^48^. Due to its immunosuppressant properties, rapamycin (sirolimus) is currently used to prevent kidney transplant rejection and to treat certain lung diseases, and several clinical trials are testing its efficiency in pathologies such as Alzheimeŕs disease and aging (Clinical Trials NCT04629495, NCT04488601). In stroke, rapamycin prevents neuronal cell death ^36, 49, 50^ and here we have observed a neuroprotective effect on apoptosis induced by OND at 24h, but not at earlier time points either after OND or tMCAo. We have also shown that rapamycin pretreatment partially prevented the impairment of microglial phagocytosis in the tMCAo model at 6h, but not in vitro in the OND model. We speculate that, in vivo, rapamycin may have indirectly improved microglial phagocytosis by reducing the tMCAo-induced ischemic damage, possibly by acting on the neurovascular unit and facilitating reperfusion ^36^. While rapamycin may not be the optimal drug to target microglia, these results suggest that preventing phagocytosis impairment in stroke models in vivo is possible.

Microglial autophagy is indeed a challenging target. We achieved autophagy inhibition using pharmacological blocking of Ulk1 or by genetic manipulation of ATG4 and Beclin1, demonstrating that basal autophagy was essential to sustain microglial survival and phagocytosis of apoptotic cells. In agreement with our results, disruption of basal autophagy has been involved in the phagocytosis of myelin ^51^ and beta amyloid deposits ^52^, in mice deficient in ATG7 and Beclin1, respectively. However, we found no significant effects in AMBRA1 heterozygous mice, suggesting that microglial autophagy may have a unique set of regulators compared to other cell types ^34^. Inhibition of the protective autophagy response mounted after OND was also detrimental for microglial survival. Unexpectedly, promoting this response with rapamycin not only did not recover phagocytosis but even had a deleterious effect on the survival of phagocytic microglia during OND. These results point to the complex regulation of autophagy in microglia, whose beneficial or detrimental effects may be time and concentration dependent. Nonetheless, microglial autophagy is a promising target to be explored. Autophagy controls the microglial inflammatory response in rodent stroke models ^53, 54^ through Annexin 1, which activates the inflammatory transcription factor NF-κB by directing its inhibitor IKK to autophagosomes for degradation ^55^. Future research will identify the target organelles or subcellular substrates that need to be recycled in microglia to maintain its health status, and whether microglial autophagy can be therapeutically exploited to support phagocytosis.

### Microglia beyond the inflammatory paradigm

For too long, the field of stroke has focused on microglial inflammatory responses with little attention to their other functions. The field is still categorizing pro- or anti-inflammatory microglia using outdated terms ^56^, such as M1 and M2 to define presumed beneficial or detrimental subtypes ^3, 54, 55, 57, 58^. In contrast, single cell RNA sequencing studies have clearly shown that microglia do not polarize to either of these categories in rodent stroke models ^59–61^. This stagnation has led to a shortage of microglial targets for clinical trials, which are to this day still largely focused on inflammation. In addition, studies in stroke patients have also oversimplified the role of human microglia by studying its “activation” in imaging studies, whereas functional studies are largely missing ^3^. Our results demonstrate that microglial phagocytosis is a promising new target in stroke, with a solid therapeutic potential of microglial phagocytosis to restore brain homeostasis that grants further exploration.

## AUTHOR CONTRIBUTIONS

SB, VST, JV, MGZ, ACG, ECZ, PRHR, DR, TU, AO, WH, CD, TF, GM, APZ and AS performed experiments and assisted with sample collection. MD, OT, PB, DS, ECS and KB provided brain samples. SB, VST, JV, MGZ, APZ and AS analyzed the results. APZ and AS supervised the study. SB, VST, APZ and AS wrote the paper, with input from all the authors. All authors reviewed and approved the manuscript.

## CONFLICTS OF INTEREST

Authors declare no competing financial interests.

## ACKNOWLEDGEMENTS

We dedicate this paper to Takashi Umekawa, who generated the HI model at the Karolinska Institute, and unfortunately passed away in 2018. This work was supported by grants from the Spanish Ministry of Science and Innovation Competitiveness MCIN/AEI/10.13039/501100011033 (https://www.ciencia.gob.es/) and ERDF “A way to make Europe” (RTI2018-099267-B-I00 and RYC-2013-12817), a Tatiana Foundation Award (P-048-FTPGB 2018)), a Basque Government Department of Education project to AS (PIBA 2020_1_0030; http://www.euskadi.eus/basque-government/department-education/) and a Basque Government Department of Economic development, Sustainability and environment (ELKARTEK KK-2020/00034; https://www.spri.eus/en/) to ECZ. SB is recipient of predoctoral fellowship from the Spanish Ministry of Economy and Competitiveness, and VST is recipient of predoctoral fellowship from the Basque Government. The funders had no role in study design, data collection and analysis, decision to publish, or preparation of the manuscript. UPV/EHU SGIker technical and human support is gratefully acknowledged.

---

**Supplementary Figure 1.**
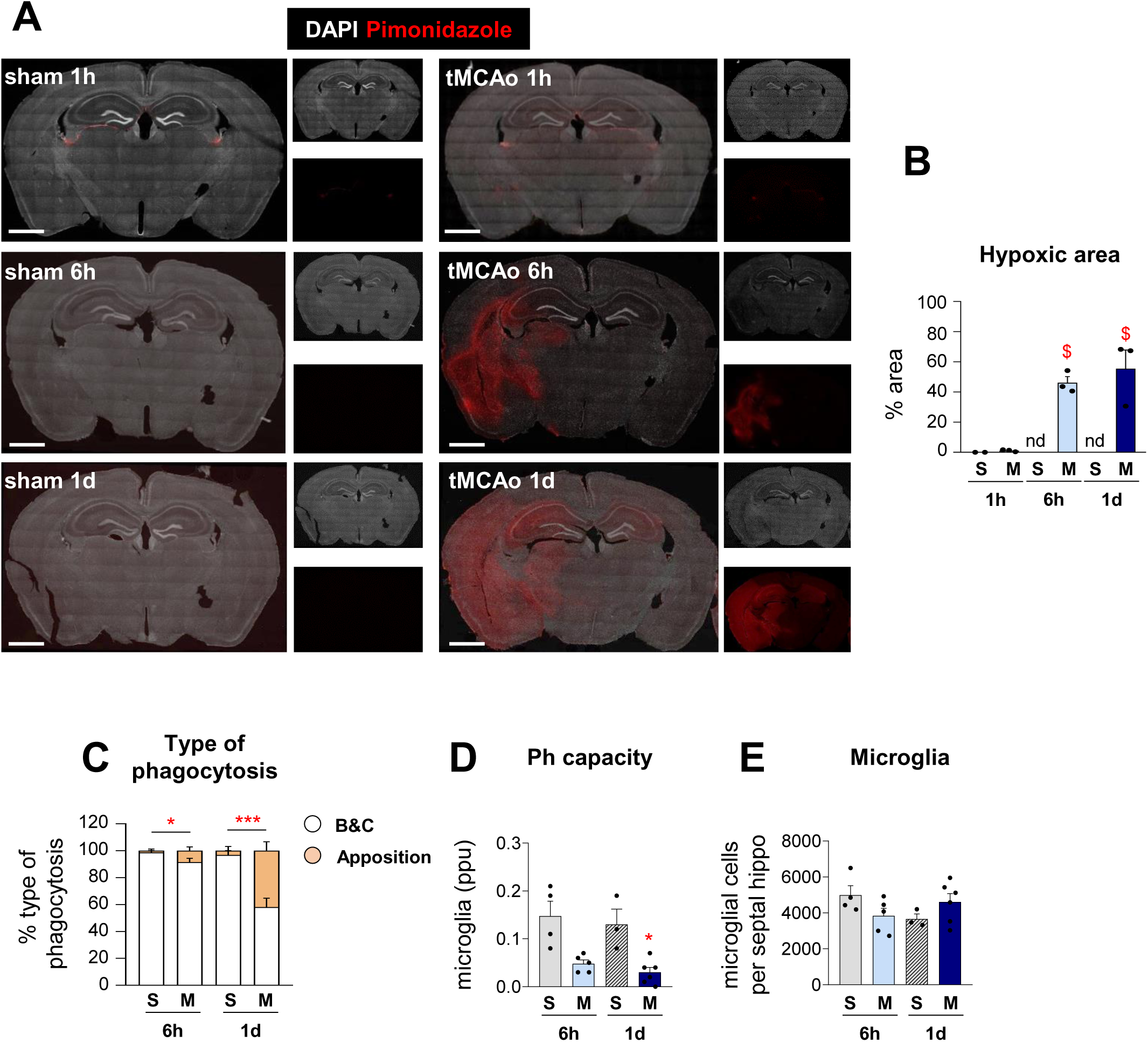
Hypoxic area and type of phagocytosis associated to tMCAo. **[A]** Representative tiled confocal image of coronal hippocampi showing cell nuclei (with DAPI, in white) and hypoxic areas labeled with the hypoxic probe pimonidazole (in red) in 2m fms-EGFP mice after sham (S) and tMCAo (M) treatment at 1h, 6h and 1d. **[B]** Percentage of hypoxic brain area determined by pimonidazole (in red) after tMCAo at 6h and 1d. The pimonidazole signal was not detected (nd) in sham animals. **[C]** Type of microglial phagocytosis (% of microglia) by “ball-and-chain” (B&C) or Apposition mechanism. **[D]** Weighted Ph capacity (% of microglia with phagocytic pouches) in sham and tMCAo. **[E]** Number of fms-EGFP^+^ microglia per septal hippocampus. Bars show mean ± SEM. In **[B]**, n=2 (sham at 1h), n=3 (sham 6h and 1d), and n=3 (tMCAo at 1h, 6h, and 1d); **[C, D, E]**, n=3 (sham at 6h), n=4 (sham at 1d), n=5 (tMCAo at 6h) and n=6 (tMCAo at 1d). Data were analyzed using 1-way ANOVA using Holm-Sidak as post hoc test [B], or 2-way ANOVA **[D, E]**. Significant interactions were found between the two factors (tMCAo treatment and time); therefore, data were split into two 1-way ANOVAs to analyze statistical differences due to the time after sham/ tMCAo at each time. Holm-Sidak was used as a post hoc test. To comply with homoscedasticity, some data were (Log_10_+1) **[C]** and/or Log_10_ **[D]** transformed. In the case that homoscedasticity was not achieved with a logarithmic transformation, data were analyzed using a Kruskal-Wallis ranks test, followed by Dunn method as a post hoc test **[C]**. (*) indicates significance compared to sham and ($) versus tMCAo at 1h. One symbol represents p<0.05 and three p<0.001. Scale bars=500μm.

**Supplementary Figure 2.**
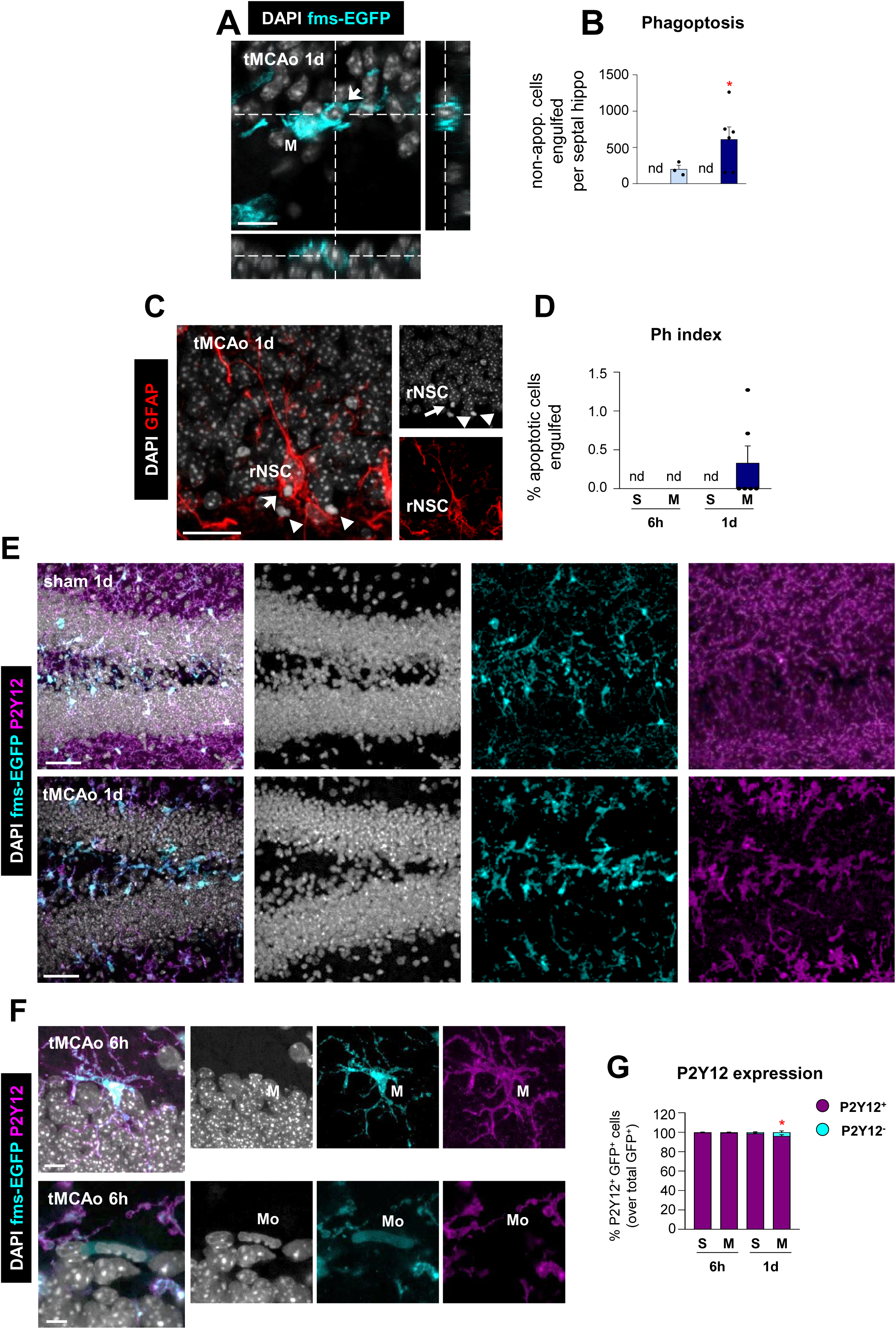
Microglial phagoptosis, rNSC phagocytosis, and P2Y12 expression. **[A]** Orthogonal projection of a non-apoptotic cell (arrow) showing its complete engulfment by microglia (fms-EGFP^+^, in cyan; M) at 1d after tMCAo. **[B]** Density of engulfed non-apoptotic cells in the septal DG in sham and tMCAo. Phagoptosis events were not detected (nd) in sham mice. **[C]** Representative confocal z-stack of an apoptotic cell (pyknotic/karyorrhetic, with DAPI, in white) phagocytosed (arrow) or not (arrowheads) by a radial neuroprogenitor (rNSC) labeled with GFAP (red) in the DG of tMCAo-treated mice at 1d. S indicates sham-mice and M tMCAo mice. **[D]** rNSC Ph index in the septal hippocampus (% of apoptotic cells engulfed by an rNSC). Apoptotic cells engulfed were not found in sham mice. nd, not-detected. **[E]** Representative confocal z-stacks of the DG of 2m fms-EGFP mice at 6h and 1d after sham and tMCAo. Cell nuclei were visualized with DAPI (in white) and microglia (fms-EGFP^+^, in cyan). Expression of P2Y12 is shown in magenta. **[F]** Representative confocal z-stack from the septal DG of a tMCAo-treated mouse at 6h showing P2Y12 expression in GFP^+^ microglia (M) and not in presumptive monocytes (Mo). **[G]** Percentage of GFP-labeled P2Y12 among the total number of GFP^+^cells in the DG at 6h and 1d after sham and tMCAo treatment. S indicates sham-mice and M tMCAo mice. Bars show mean ± SEM. In **[B and D]**, n=5 (tMCAo 6h) and n=6 (tMCAo 1d); **[G]**, n=3 per group (sham/tMCAo at 6h and 1d). Data were analyzed using Student s t test **[B]**, or 2-way ANOVA **[G]**. Significant interactions were found only across the time. Data were split into two 1-way ANOVAs to analyze statistical differences due to the time after sham/ tMCAo at each time. Holm-Sidak was used as a post hoc test. (^$^) represents p<0.05 compared to tMCAo at 6h. Scale bars=10μm, z=11.2μm **[A]**; 20μm, z=10μm **[C]**, 50μm, z= 20μm **[E];** 10μm, z=9.8μm (tMCAo at 6h), 6.3μm (tMCAo at 1d) **[F]**.

**Supplementary Figure 3.**
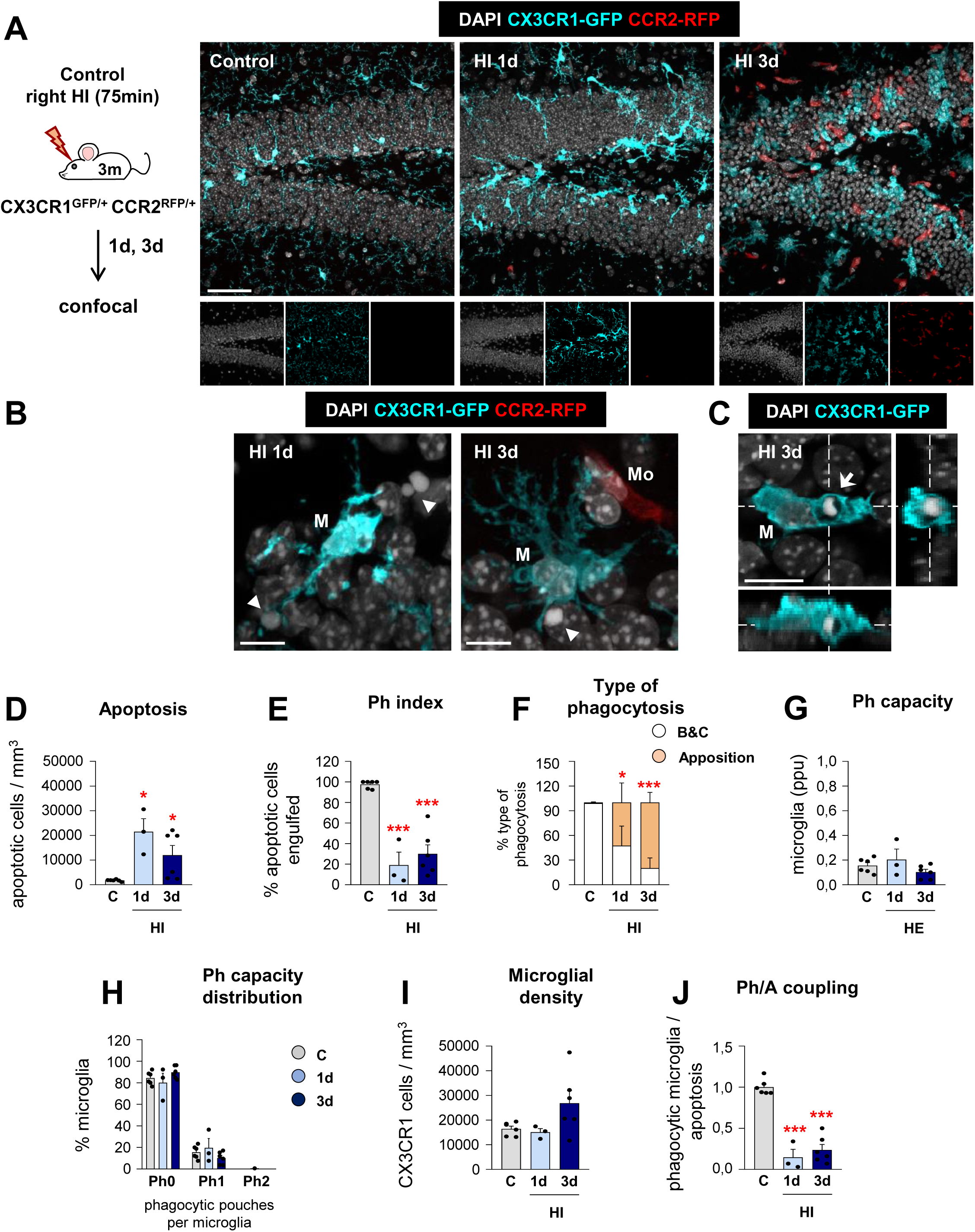
Microglial phagocytosis is impaired during HI. **[A]** Experimental design and representative confocal z-stack of the DG of 3 month-old (3m) CX3CR1-GFP CCR2-RFP mice at 1 and 3d under hypoxia-ischemia (HI). Cell nuclei were visualized with DAPI (in white), microglia (CX3CR1-GFP^+^, in cyan) and monocytes (CCR2-RFP^+^, in red). **[B]** Representative confocal z-stack from the DG of a HI-treated mouse at 1 and 3d showing apoptotic cells (arrowheads) non-phagocytosed by microglia (CX3CR1-GFP^+^, in cyan; M), close to monocytes (CCR2-RFP^+^, in red; Mo). **[C]** Orthogonal projection of a confocal z-stack from the septal DG of a HI-treated mouse at 3d showing an apoptotic cell (arrow) phagocytosed by microglia (CX3CR1-GFP^+^, in cyan; M). **[D]** Density of apoptotic cells (cells/mm^3^) in the septal DG under control and HI treatment. **[E]** Ph index in the septal DG (in % of apoptotic cells engulfed by microglia). **[F]** Type of microglial phagocytosis (in % of microglia) by “ball-and-chain” (B&C) or “Apposition” (Appo) mechanism. **[G]** Weighted Ph capacity under control and HI conditions. **[H]** Histogram showing the Ph capacity of microglia (% of microglia with pouches). **[I]** Density of CX3CR1-GFP^+^ microglia(cells/mm^3^) in the septal DG. **[J]** Ph/A coupling (in fold change) in the septal DG. Bars represent mean ± SEM. n=5 (control), n=3 (at 1d) and n=6 (at 3d). Data was analyzed by 1-way ANOVA, using Holm-Sidak as post hoc test. To comply with homoscedasticity, some data were Log_10_ **[D, H]** and/or Log_10_+1 **[G]** transformed. In the case that homoscedasticity was not achieved with a logarithmic transformation data were analyzed using a Kruskal-Wallis ranks test, followed by Dunn method as a post hoc test **[D, G]**. One (*) symbol indicates p<0.05, and three p<0.001 (vs control). Scale bars=50μm, z=16μm **[A]**; 20μm, z=15μm **[B]**; 10μm, z=14μm **[C]**.

**Supplementary Figure 4.**
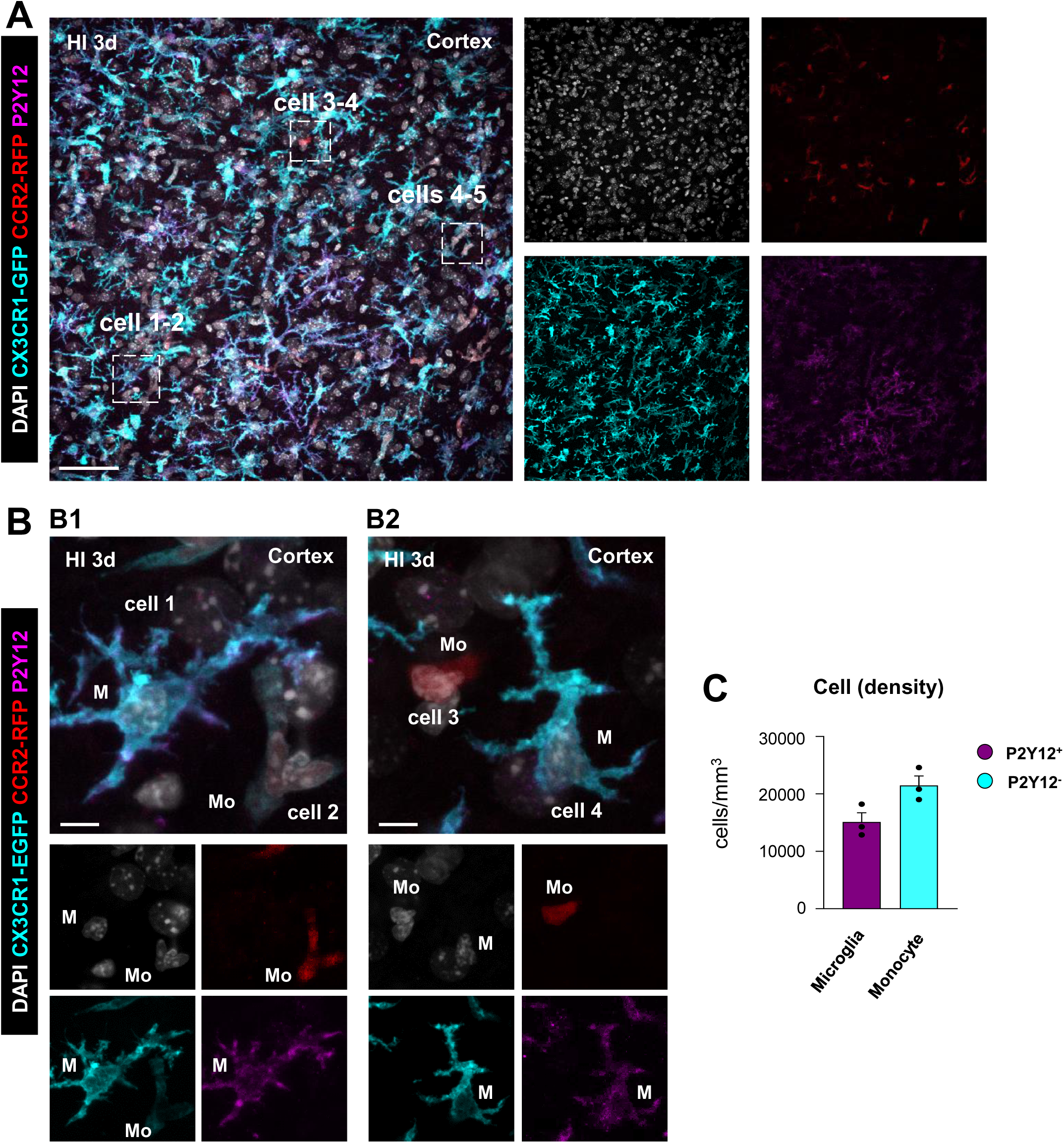
P2Y12 expression discriminates microglia from peripheral monocytes in the adult brain after HI. **[A]** Representative confocal z-stack of the DG of 3 month (3m) CX3CR1-GFP CCR2-RFP mice at 3d under hypoxia-ischemia conditions (HI). Cell nuclei were visualized with DAPI (in white), microglia (CX3CR1-GFP^+^, in cyan), and monocytes (CCR2-RFP^+^, in red). Expression of P2Y12 is shown in magenta. **[B]** Representative confocal z-stack from the septal DG of a HI-treated mouse at 3d showing a P2Y12 expression (in magenta) in CX3CR1-GFP^+^ microglia (cells 1 and 4, M) and not in CCR2-RFP^+^ monocytes(cells 2, 3, Mo) **[B3]**. **[C]** Density of microglia (GFP^+^ RFP^-^) and monocytes (GFP^-^ RFP^+^ and GFP^+^ RFP^+^ cells/mm^3^) expressing P2Y12 in the cortex at 3d after HI treatment. Bars represent mean ± SEM. The expression of P2Y12R on microglia and monocytes **[C]** was analyzed using Student’s t test. n=3 mice per group. Scale bars=50μm, z=12μm **[A]**; 10μm **[B]**, 10μm **[B1-B2]**.

**Supplementary Figure 5.**
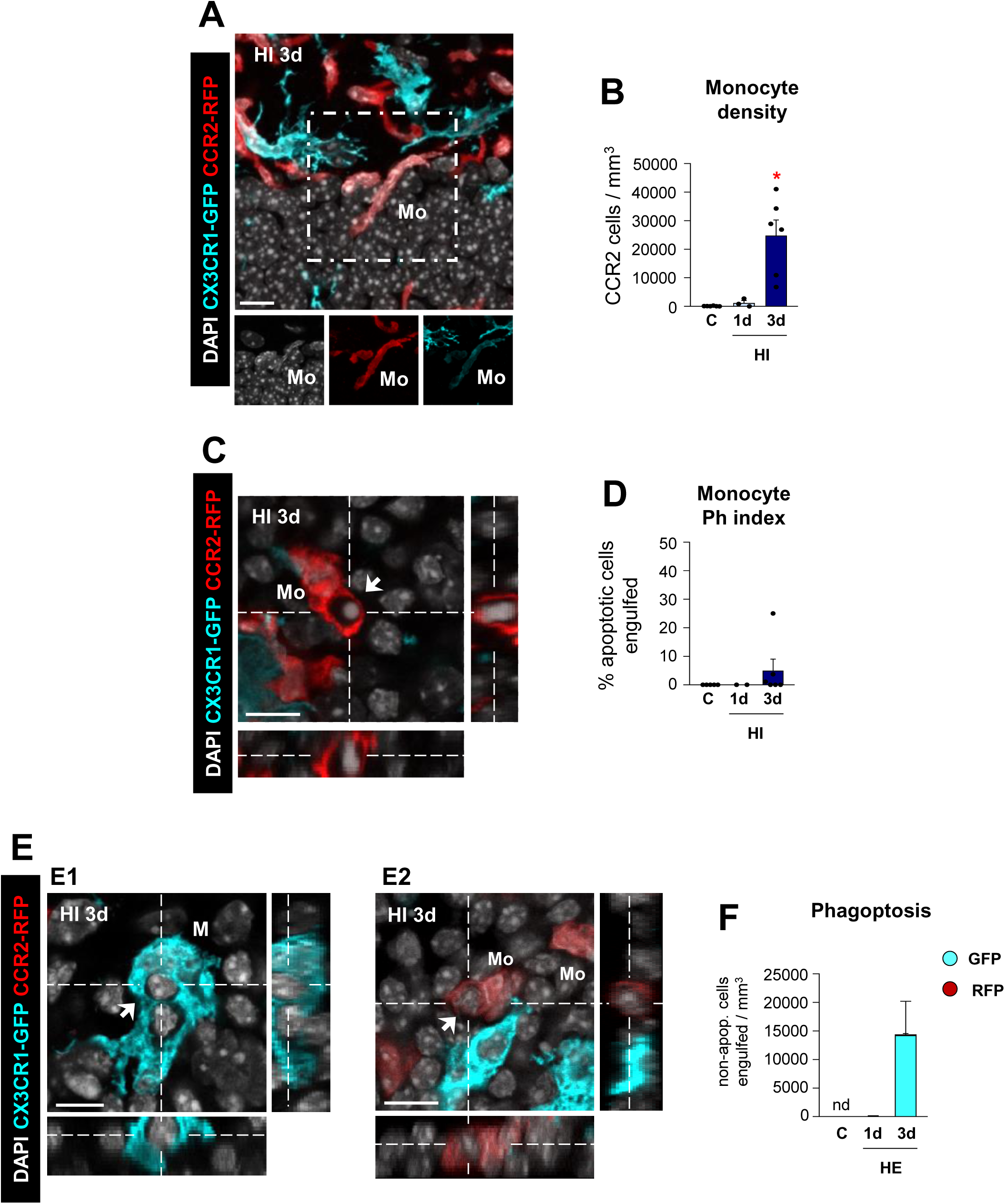
Microglial phagocytosis impairment induced by HI is not compensated by monocyte phagocytosis. **[A]** Representative confocal z-stack from the DG of a CCR2-RFP^+^ cell (in red; Mo) expressing GFP signal (in cyan) at 3d after HI. **[B]** Density of monocytes (in CCR2-RFP^+^ cells/mm^3^) in the septal DG under control and HI conditions. **[C]** Orthogonal projection of a confocal z-stack from the DG of a HI-treated mouse at 3d showing an apoptotic cell (arrow) phagocytosed by a monocyte (CCR2-RFP+, in red; Mo). **[D]** Monocyte Ph index in the septal DG (% of apoptotic cells engulfed by monocytes). **[E]** Orthogonal projection of non-apoptotic cell (arrow) showing its complete engulfment by microglia (CX3CR1-GFP^+^, in cyan; M) **[E1]** or peripheral monocytes (CCR2-RFP^+^, in light red; Mo)**[E2]** at 3d after HI. **[F]** Density of engulfed non-apoptotic cells in the septal DG (in cells/mm^3^) in control and HI treatment. Phagoptosis was not detected (nd) in control mice. Bars represent mean ± SEM. In **[B, D, F]**, n=5 (control), n=3 (at 1d) and n=6 (at 3d). Data was analyzed by 1-way ANOVA using Holm-Sidak as post hoc test **[B, D]**, or by Student s t test **[F]**. To comply with homoscedasticity, some data were Log_10_+1 **[B]** transformed. * indicates p<0.05 (vs control). Scale bars=50μm, z=15μm **[A]**; 20μm, z=12μm **[C]**; 20μm, z=10.5μm **[E]**.

**Supplementary Figure 6.**
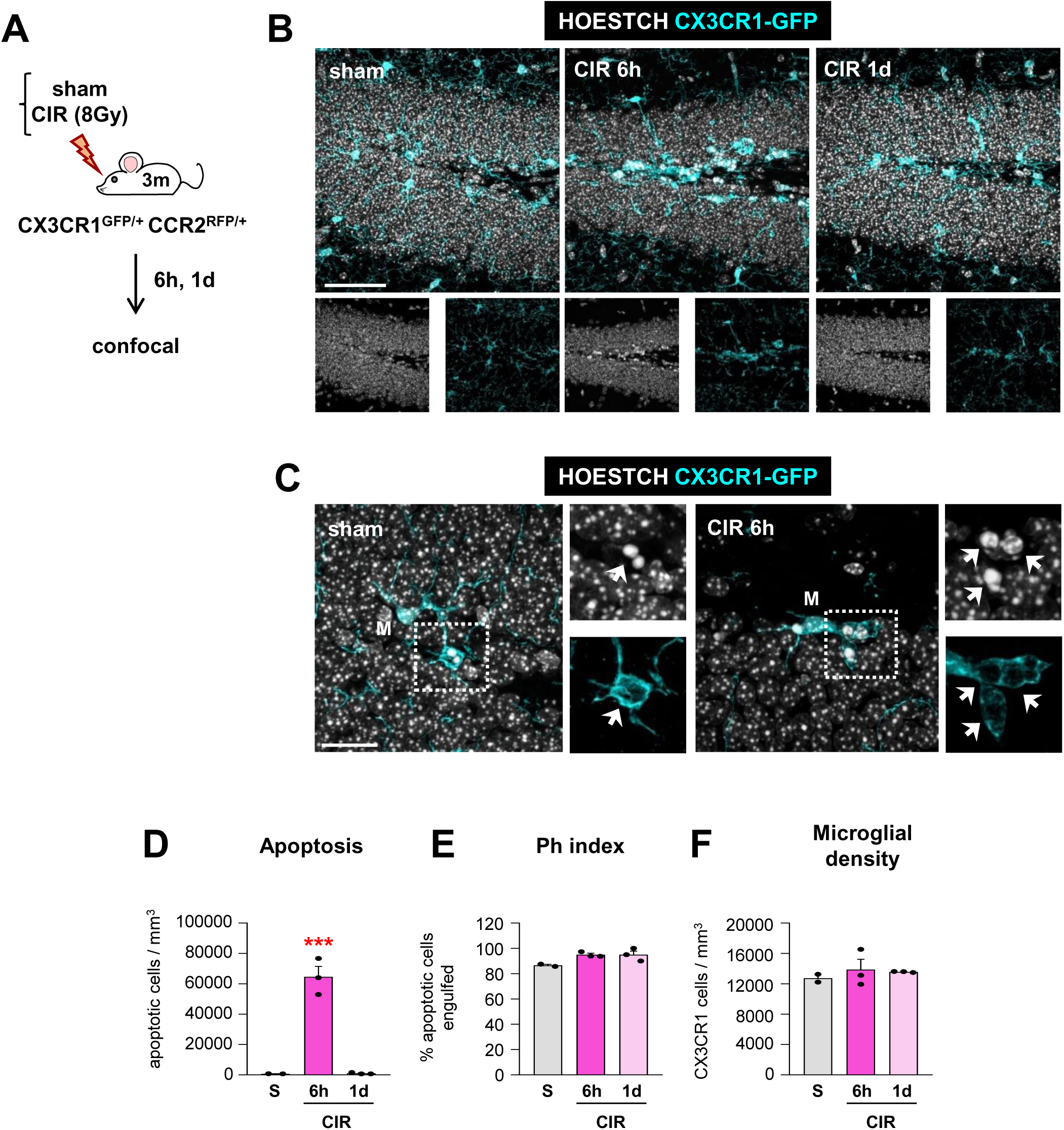
Microglial phagocytosis increases after cranial irradiation (CIR) exposure. **[A, B]** Experimental design and representative confocal z-stacks of the DG of 3m CX3CR1-GFP/CCR2-RFP mice at 6h and 1d after cranial irradiation (CIR, 8Gy). Apoptotic nuclei were detected by pyknosis/karyorrhexis (in white, Hoestch), microglia and blood-derived macrophages by the transgenic expression of CX3CR1-GFP (in cyan) and CCR2-RFP (in red), respectively. **[C]** Representative confocal z-stack of apoptotic cells (pyknotic/karyorrhectic, Hoescht, in white, arrow) phagocytosed by microglia (CX3CR1-GFP^+^, in cyan; M) in the septal DG of sham and CIR-treated mice. **[D]** Density of apoptotic cells (cells/mm^3^) in the septal DG. **[E]** Ph index (% of apoptotic cells engulfed by microglia) in the septal DG. **[F]** Density of CX3CR1^+^ microglia (cells/mm^3^) in the septal DG. Bars represent mean ± SEM. In **[C, D, E]**, n=2 (sham) and n=3 (at 6h and 1d). Data was analyzed by 1-way-ANOVA, using Holm-Sidak as a post hoc test. To comply with homoscedasticity some data were Log_10_ transformed **[D]**. *** indicates p<0.001 (vs CIR at 1d). Scale bar=50µm, z=21µm (sham, 6h), z=17.5µm (1d) **[A]**; 20µm, z=13.3µm (sham), z=18.9µm (6h) **[B]**.

**Supplementary Figure 7.**
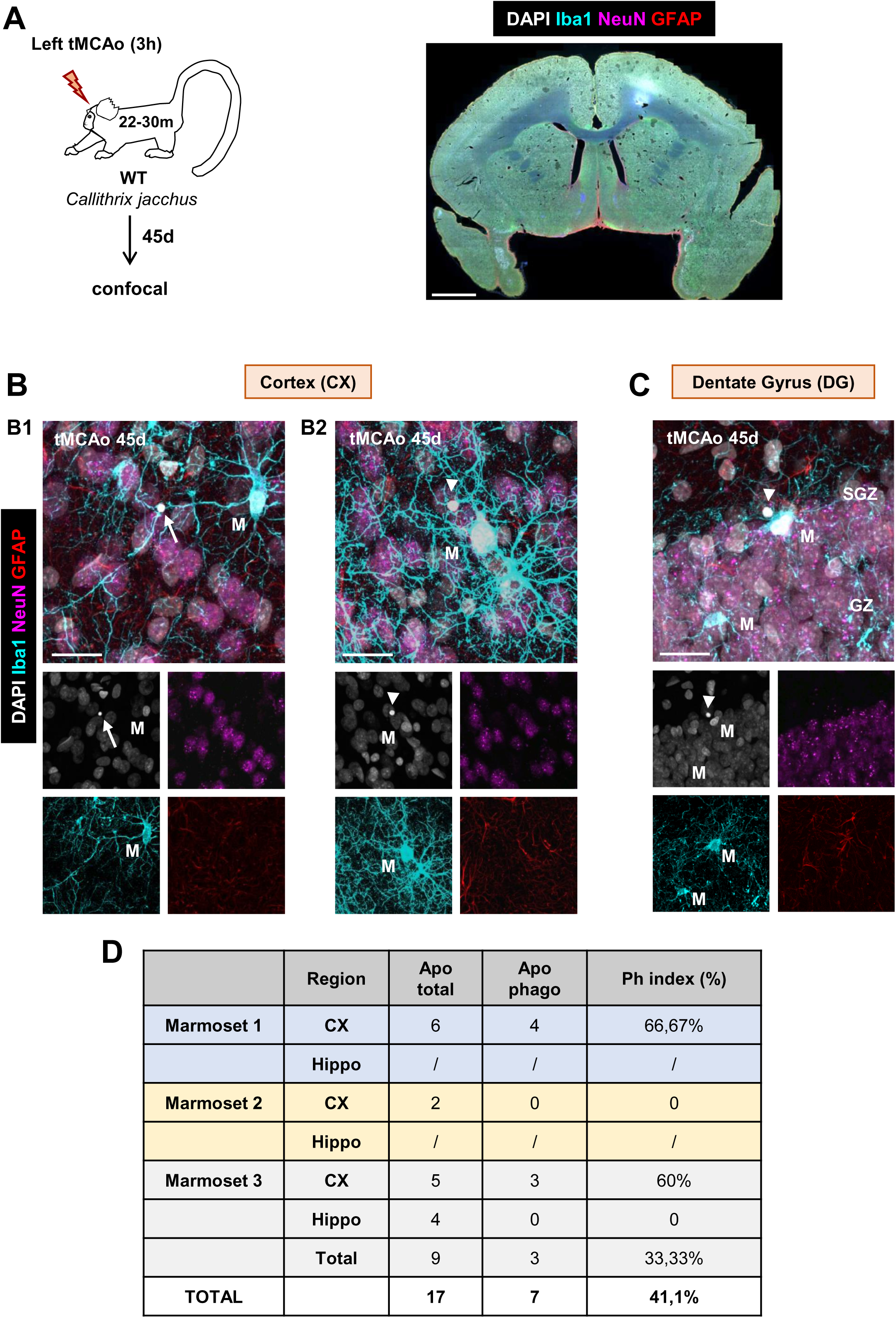
Microglial phagocytosis after tMCAo in *Callithrix jacchus*. **[A]** Experimental design of tMCAo in *Callithrix jacchus* and low magnification epifluorescent image of common marmoset brain showing nuclei (DAPI, white), microglia (Iba1, cyan), neurons (NeuN, magenta), and astroyctes (GFAP, red). **[B. C]** Representative confocal z-stacks of the cortical regions (**B**) and hippocampus (**C**) of marmosets at 45d after tMCAo showing phagocytosed (arrow) and non-phagocytosed (arrowheads) apoptotic cells. M, microglia **[D]** Table summarizing the number of apoptotic cells (total and phagocytosed by microglia) in the three marmosets analyzed. Scale bars=20μm, z=19.6μm **[B1, B2]**; 20μm, z=23.8μm **[C].**

**Supplementary Figure 8.**
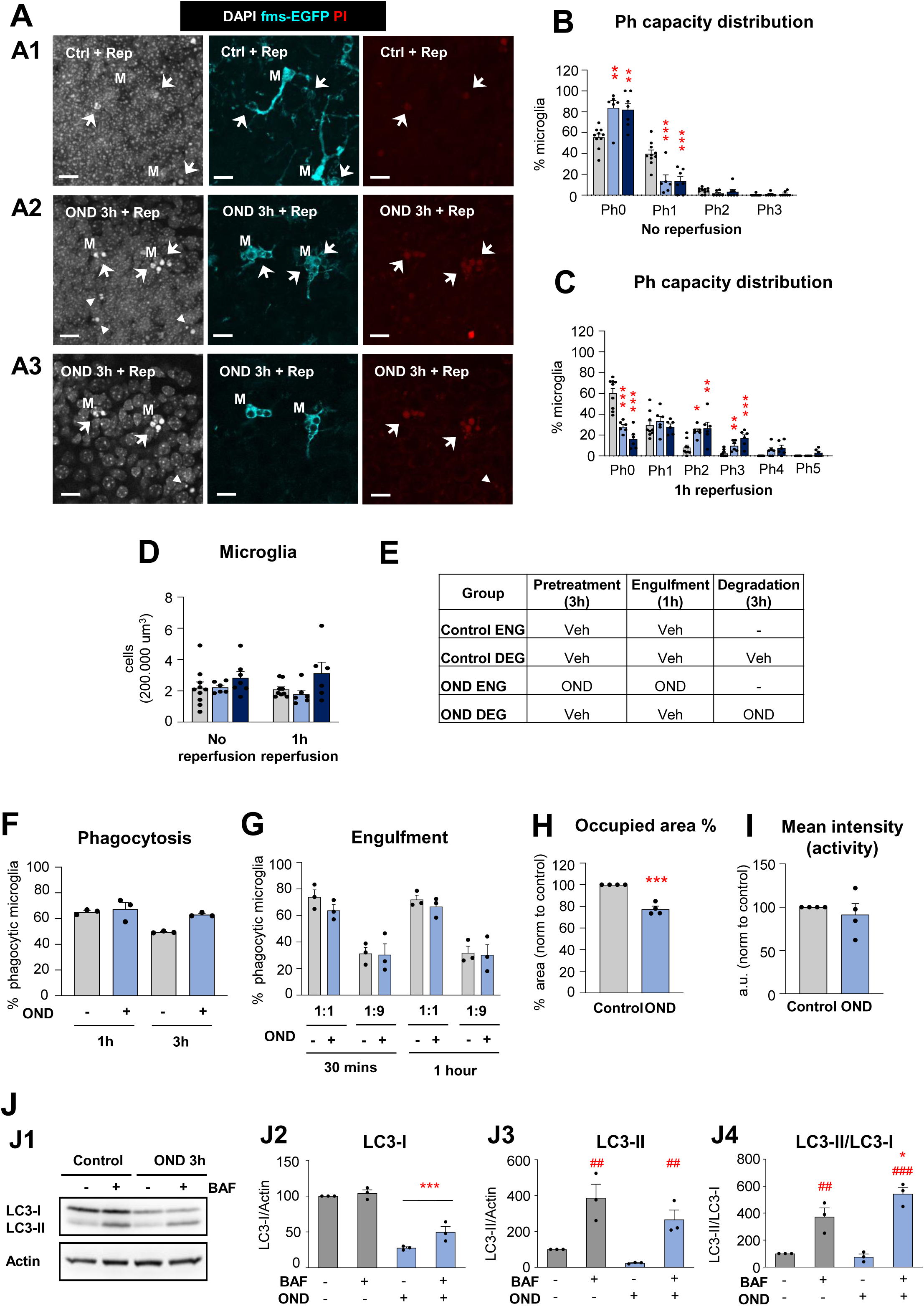
Microglial phagocytosis under OND. **[A, B, C, D]** Hippocampal organotypic slices were exposed to OND during 3 and 6h with and without 1h reperfusion. **[A]** Single channel confocal z-stacks images of those shown in **Figure 2A3** of the DG in control (**A1**) after OND and reperfusion (**A2**; single plane image is shown in **A3**). Normal or apoptotic (pyknotic/karyorrhectic) nuclear morphology was visualized with DAPI (white) and microglia by transgenic expression of fms-EGFP (cyan). Under reperfusion conditions, membrane permeability (characteristic of necrotic cells) was observed with PI (red). The images show primary apoptotic cells (pyknotic/karyorrhectic) or secondary necrotic cells (pyknotic/karyorrhectic, PI^+^) engulfed (arrows) or not-engulfed (arrowheads) by microglia (M) (EGFP^+^). **[B, C]** Ph capacity histogram after OND in non-reperfused **[B]** and reperfused **[C]** conditions. **[D]** Number of microglia in 200.000 µm^3^ of the DG. **[E]** Summary of the experimental groups in **Figure 2 J, K, L.** ENG, engulfment (1h); DEG, degradation (3h). **[F]** Percentage of phagocytic microglia 1 and 3h after the addition of apoptotic cells (raw data). **[G]** Percentage of phagocytic microglia after the addition of different apoptotic SH-SY5Y vampire neuron to microglia ratios (1:1 and 1:9) after 30 min and 1h of engulfment under OND conditions. **[H]** Percentage of the area occupied by lysosomes, referred to control values. **[I]** Mean intensity (representative of lysosomal activity) represented in arbitrary units and referred to control values. **[J]** BV2 cells were exposed to OND for 3 hours in the presence and absence of bafilomycin-A1 (BAF, 100 nM, 3 hours) to assess autophagy flux by LC3 turnover assay. Delipidated (∼1 KDa) and lipidated (∼17 KDa) LC3 levels were analyzed by western blot. Beta-actin (∼42 KDa) was used as loading control. Representative blots showing LC3-I, LC3-II and actin bands **[J1]**, LC3-I levels normalized to actin **[J2]**, LC3-II levels normalized to actin **[J3]**, LC3-II levels normalized to LC3-I levels **[J4]**. Bars show mean ± SEM. n = 6-10 mice per group **[A, B, C, D]**, n=3 independent experiments **[E-G]**, n=4 independent experiments **[H, I]**, n=4 independent experiments **[J]**. Data was analyzed by one-way ANOVA (factor: number of pouches) followed by Holm-Sidak post hoc tests **[B, C]**, two-way ANOVA followed by Holm-Sidak post hoc tests **[D]**, Student’s t-test **[H]** and by two-way ANOVA followed by Holm-Sidak post hoc test **[J].** When an interaction between factors was found, one-way ANOVA was performed followed by Holm-Sidak post hoc tests **[J4]**. (* and #) represent significance between control and OND or between bafilomycin and no bafilomycin respectively: one-symbol represents p<0.05; two-symbols represent p<0.01; three-symbols represent p<0.001 (OND vs control).

**Supplementary Figure 9.**
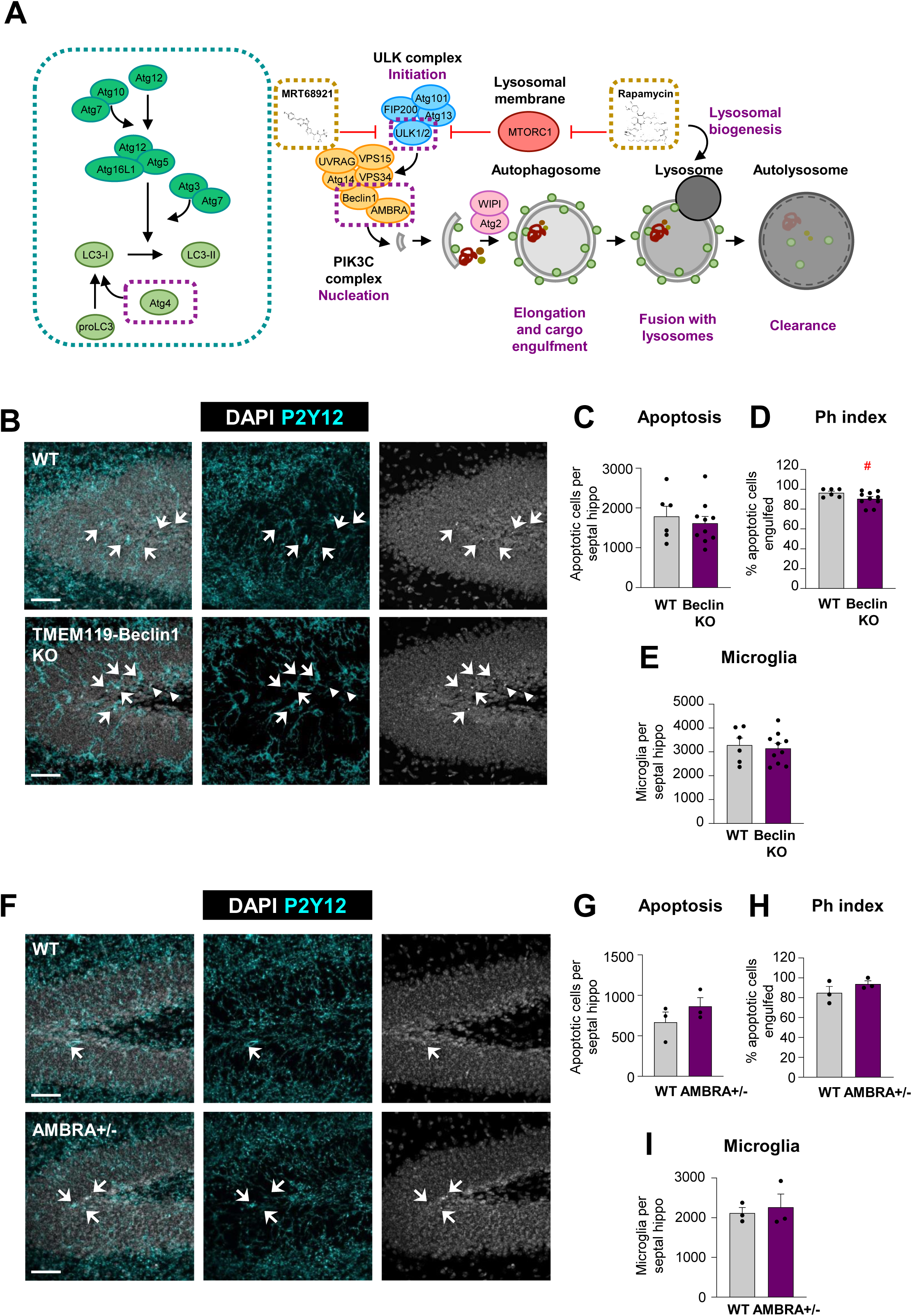
Microglial phagocytosis in mice deficient in autophagy genes. **[A]** Graphical representation of the general autophagy process and its main proteins. Target proteins used for the KO models are indicated with a purple dotted box. MRT68921 and rapamycin are represented next to their pharmacological targets in a yellow dotted box. **[B]** Representative confocal images of the DG of wild-type (WT) and TMEM-Beclin knock-out (Beclin KO) mice. Nuclei were visualized with DAPI (white) and microglia was stained for P2Y12 (cyan). Arrows point to engulfed and arrowheads to non-engulfed apoptotic cells. **[C]** Number of apoptotic cells per septal hippocampus in WT and Beclin KO mice. **[D]** Ph index in the septal hippocampus (% of apoptotic cells engulfed by microglia). **[E]** Number of microglial cells per septal hippocampus in WT and Beclin KO mice. **[F]** Representative confocal images of the DG of wild-type (WT) and heterozygous AMBRA 1 (AMBRA +/-) mice. Nuclei were visualized with DAPI (white) and microglia was stained for P2Y12 (cyan). Arrows point to engulfed apoptotic cells. **[G]** Number of apoptotic cells per septal hippocampus in WT and AMBRA +/- mice. **[H]** Ph index in the septal hippocampus (% of apoptotic cells engulfed by microglia). **[I]** Number of microglial cells per septal hippocampus in WT and AMBRA +/- mice. Bars show mean ± SEM. n=6-10 animals per group **[C-E]**; n=3 per group **[G-I].** Data was analyzed by Student’s t-test. # represents p=0.080. Scale bars=50µm **[B, F],** z=26.6µm **(WT)**, 21µm (TMEM119-Beclin1 KO) **[B]**, 16.1µm **[F]**.

**Supplementary Figure 10.**
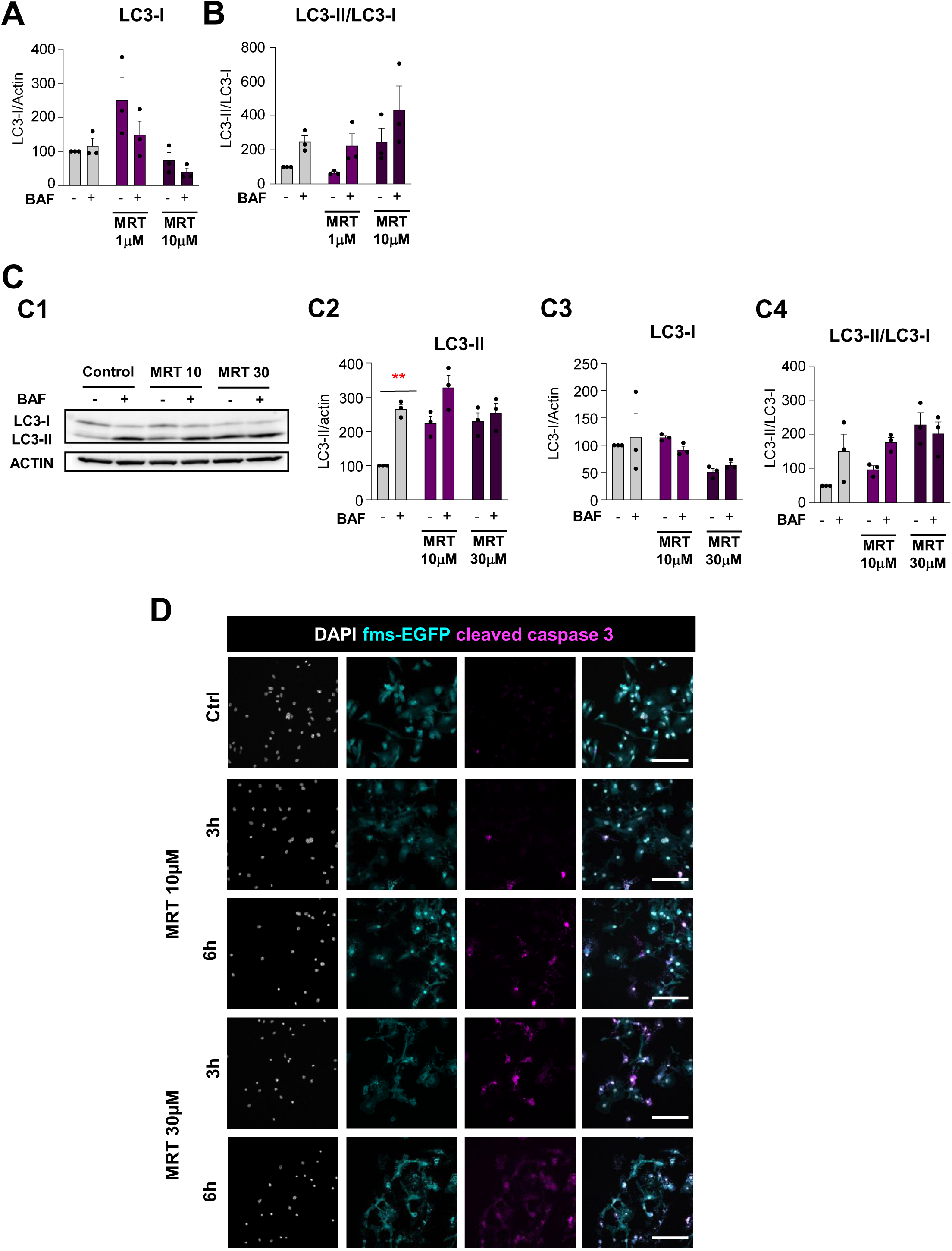
Effects of MRT68921 on microglia. **[A]** LC3-I quantification (referred to actin) in primary microglia after the addition of 1 and 10μM for 6h in the presence and absence of the lysosomal inhibitor bafilomycin (BAF, 100 nM). **[B]** LC3-II quantification (referred to LC3-I) in primary microglia after the addition of 1 and 10μM for 6h in the presence and absence of the lysosomal inhibitor bafilomycin (BAF, 100 nM). **[C] (C1)** Representative blot showing relative levels of LC3-I and LC3-II in primary microglia 3h after administration of the Ulk1/2 inhibitor MRT68921 (10 and 30μM). **(C2**, **3)** Quantification of the LC3-II and LC3-I levels referred to actin and **(C4)** LC3-II levels referred to LC3-I after 10 and 30μM MRT68921 in the presence and absence of the lysosomal inhibitor, bafilomycin A (BAF, 100 nM). These data are reprinted with permission from Frontiers in Immunology ^31^. **[D]** Representative confocal images of the MRT68921 dose-time course in primary microglia. Healthy and apoptotic nuclei (pyknotic/karyorrhectic) were assessed with DAPI (white), microglia (fms-EGFP, cyan) and apoptosis was assessed with activated caspase 3 (act-casp3^+^, magenta). Bars show mean ± SEM. n=3 **[A, B]**, n=3 **[C]** Data was analyzed by two-way ANOVA followed by Holm-Sidak post hoc test **[C]** (*) represents significance between bafilomycin and no bafilomycin, respectively: two symbols represent p<0.01. Scale bar=50μm, z=8.5μm.

**Supplementary Figure 11.**
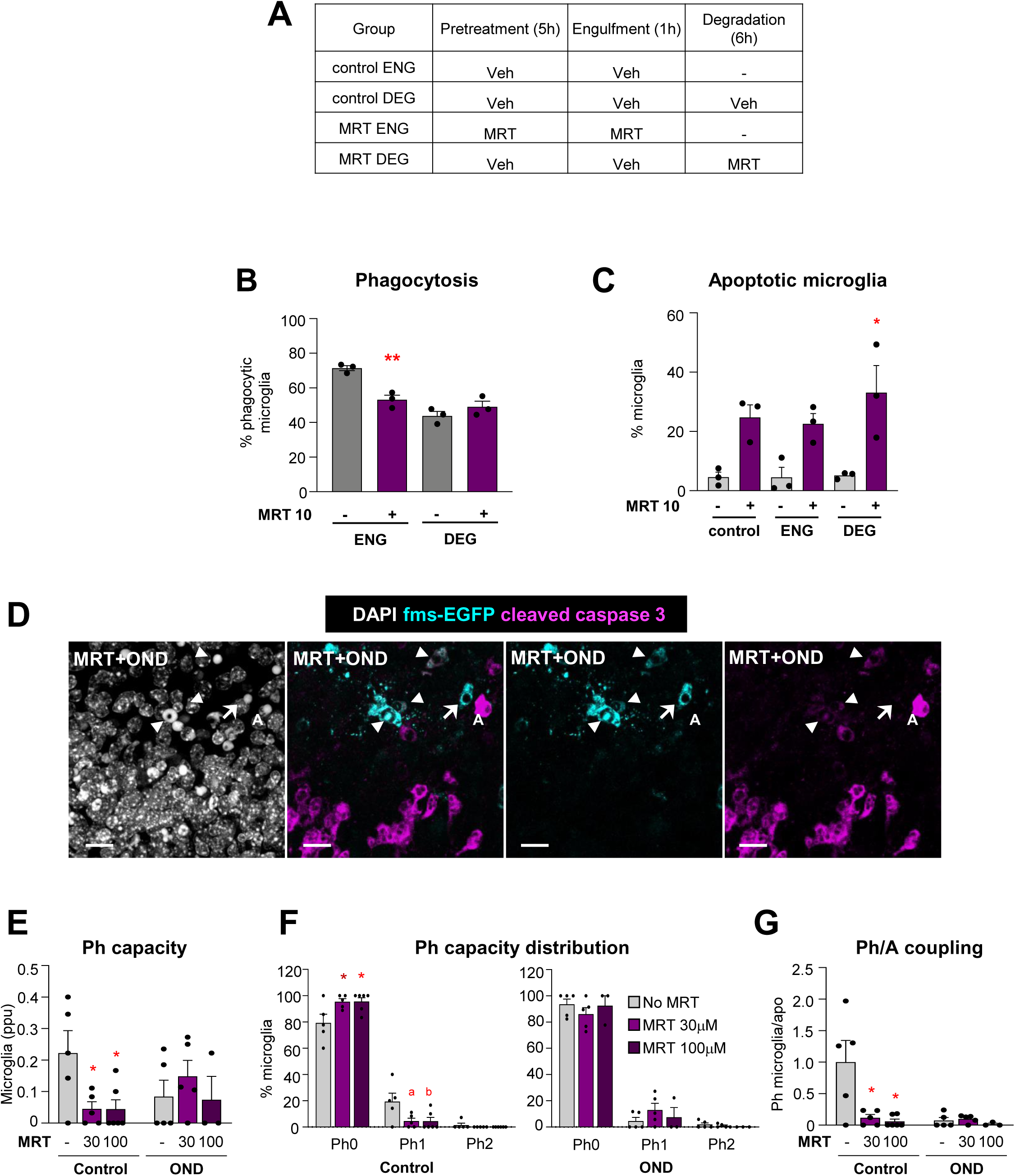
Basal autophagy is essential to maintain microglial function and survival. [**A**] Summary of the experimental groups in **Figure 4 I, J, K**. [**B**] Percentage of phagocytic microglia after 1h (engulfment) and 6h (degradation) after the addition of apoptotic cells in microglia treated with the Ullk1/2 inhibitor MRT68921 (10 μM; raw data). [**C**] Percentage of pyknotic/karyorrhectic microglia in the control and phagocytic microglia during the engulfment and degradation steps, in the presence of 10 μM MRT68921. **[D]** Representative high-magnification confocal images of apoptotic microglia after 3h treatment with MRT68921 (100 µM) and exposure to OND. Normal or apoptotic (pyknotic/karyorrhectic) nuclear morphology was visualized with DAPI (white), apoptosis was confirmed by activated caspase-3 staining (red), and microglia by the transgenic expression of fms-EGFP (cyan). Arrowheads, apoptotic microglia (fms-EGFP^+^, act-casp3^+^); arrow, phagocytic microglia (fms-EGFP^+^, act-casp3^-^); A, apoptotic cell (act-casp3^+^ with pyknotic/karyorrhectic nuclear morphology). **[E, F, G]** Hippocampal organotypic slices were treated with MRT68921 (30 and 100 µm) for 3h in the presence and absence of OND. **[E]** Weighted Ph capacity (number of phagocytic pouches filled with apoptotic nuclei per microglia) in parts per unit (ppu). **[F]** Ph capacity histogram in control and OND conditions. **[G]** Ph/A coupling in fold-change (net phagocytosis with respect to total levels of apoptosis). Bars represent the mean ± SEM. n=3 independent experiments **[B, C]**, n=3-6 mice per experimental condition **[E-G]**. Some data was transformed to comply with homoscedasticity using Log_10_ **[E]** or square root **[G]**. Data was analyzed by 1-way ANOVA followed by Tukeýs multiple comparisons in case significant differences were identified [**B**], by two-way ANOVA followed by one-way ANOVA (factor: MRT treatment) when an interaction was revealed in data split in control and OND conditions and Holm-Sidak post hoc tests **[E,G]** and one-way ANOVA (factor: number of pouches) followed by Holm-Sidak post hoc tests **[F].** (* and #) represent significance vs the control group or between MRT-treated and untreated groups: one symbol represents p<0.05 and two symbols represent p<0.01.; (a) represents p=0.06 (MRT30 vs untreated), (b) represents p=0.07 (MRT100 vs untreated). Scale bars=50μm, z=8.5 µm [**B, C**]; 15µm, z=12.6 µm **[D]**.

**Supplementary Figure 12.**
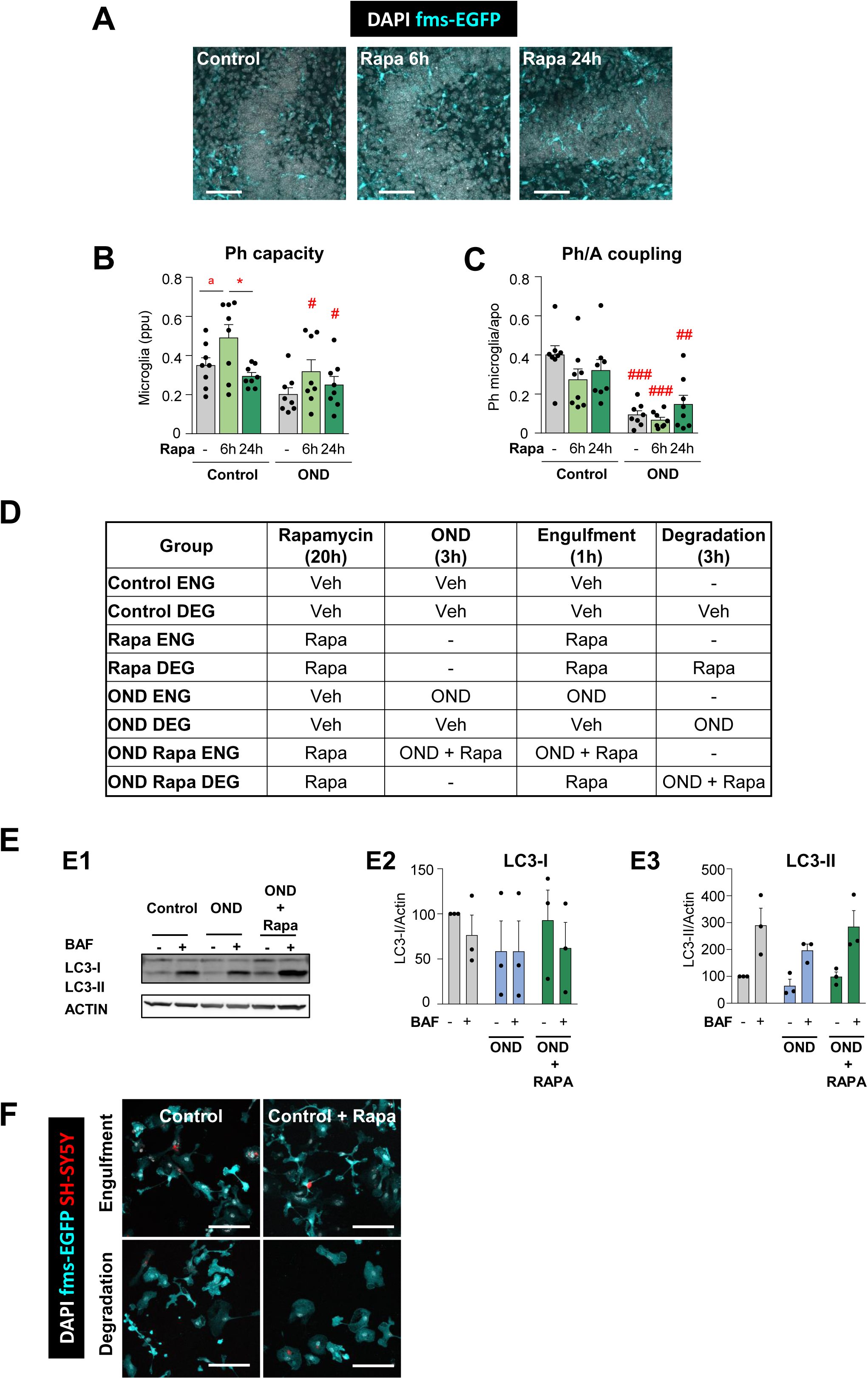
Rapamycin does not revert the phagocytic impairment in vitro. **[A]** Representative confocal images of the DG after rapamycin (200 nM) treatment for 6 and 24h under control conditions. Normal or apoptotic (pyknotic/karyorrhectic) nuclear morphology was visualized with DAPI (white) and microglia by the transgenic expression of fms-EGFP (cyan). **[B, C]** Quantification of the **[B]** weighted Ph capacity (number of phagocytic pouches filled with apoptotic nuclei per microglia) in parts per unit (ppu), and **[C]** Ph/A coupling, in fold-change (net phagocytosis with respect to total levels of apoptosis) after treatment of hippocampal organotypic slices with rapamycin (200nM) for 6 and 24h in control and OND conditions. **[D]** Summary of the experimental groups in **Figure 5 K, L, M**. **[E]** Representative blot showing relative levels of LC3-I and LC3-II in primary microglia after 21h of rapamycin pretreatment followed by 3h of OND **[E1]**. Quantification of the LC3-I and LC3-II levels (referred to actin) after 21h of rapamycin pretreatment followed by 3h of OND **[E2**, **E3]**. **[F]** Control images corresponding to **Figure 5J**. Representative confocal images of control microglia (GFP, cyan) after 1 (engulfment) and 3h (degradation) of apoptotic SH-SY5Y vampire neurons (red) after rapamycin pretreatment. Bars represent mean ± SEM. n=8 mice per group **[B, C]**, n=3-4 independent experiments **[E]**. Some data was transformed using a square root to comply with homoscedasticity **[C]**. Data was analyzed by two-way ANOVA followed by Holm-Sidak post hoc tests **[B, C]** .(* and ^#^) represent significance between control vs rapamycin, and control vs OND, respectively **[B, C]**; (*) represent significance between degradation under OND and rapamycin compared to OND, OND with rapamycin and engulfment under OND and rapamycin: one symbol represents p<0.05, two symbols represent p<0.01 and three symbols represent p<0.001; (a) represents p=0.079 (rapamycin 6h vs control). Scale bars=50µm, z=11.2µm **[A]**.

## MATERIALS AND METHODS

### Mice

All experiments were performed in adult (2 month old) fms-EGFP (MacGreen) mice, except where indicated, in which microglia constitutively express the green fluorescent reporter under the expression of the c-fms promoter ^17, 63^. Two-photon microscopy experiments were performed in CX3CR1^GFP/+^ where microglia express GFP ^64^. Brain tissue from CX3CR1^GFP/+^ CCR2^RFP/+^ mice was provided by K. Blomgren (Karolinska Institute, Sweden), in which peripheral monocytes express the red fluorescent protein RFP ^65^. Brain tissue from constitutive ATG4B KO was provided by G. Mariño, Universidad de Oviedo, Spain ^66^; AMBRA1^+/-^ heterozygous mice by P. Boya, Centro de Investigaciones Biológicas CIB, Spain ^33^; and tamoxifen inducible KO of Beclin1 under the TMEM119-CreERT2 by D. Schafer, University of Massachussetts, USA. All animals used were on a C57BL/6 background except ATG4B KO mice and AMBRA1+/- heterozygous mice, which were on a mixed C57BI6/129 Sv and CD1 background, respectively. Mice were housed in 12:12h light cycle with ad libitum access to food and water. Both males and females were used and pooled together, unless otherwise noted. All procedures followed the European Directive 2010/63/EU, NIH guidelines and were approved by the Ethics Committees of the University of the Basque Country EHU/UPV (Leioa, Spain; CEBA/205/2011, CEBA/206/2011, CEIAB/82/2011, CEIAB/105/2012) and Karolinska Institute (Stockholm, Sweden; protocol number N249/13).

### Non-Human Primates

Brain samples from cynomolgus macaque (*Macaca fascicularis*) and common marmoset (*Callithrix jacchus*) under transient Middle Cerebral Artery occlusion were provided by Emmanuelle Canet-Soulas (University of Lyon, France) ^22^ and Omar Touzani (University of Caen, France)^25^, respectively. Animals were housed in a 12h light-dark cycle. All procedures were approved by European Directives and approved by the Regional Ethics Committee. For tMCAo in non-human primates, 7 years (7y) cynomolgus macaque and 2-2.5y common marmosets were anaesthetized with sevoflurane and isofluorane, respectively, as previously described ^25, 62^, and MCA was occluded for 110min in macaque and 3h in marmosets. Animals were sacrificed at 30d and 45d after tMCAo, respectively.

### Primary microglia

Primary microglia was obtained as previously ^67^ described. Briefly, postnatal day 0-1 (P0-P1) fms-EGFP or wildtype (WT) mice were sacrificed by decapitation and brains were extracted. Meninges were removed in Hank’s balanced salt solution (HBSS, Gibco) under a binocular magnifier and the cerebellum and olfactory bulbs were removed. The remaining brain was manually chopped and enzymatically digested in the presence of papain (20U/mL, Sigma), a cysteine protease enzyme extracted from the papaya latex, and desoxyribonuclease I (DNAse I, 150U/μL, Gibco) for 10-15 minutes at 37°C after which the remaining fragments were mechanically homogenized by gentle pipetting. The resulting cell suspension was filtered through a 40μm polypropylene cell strainer (Fisher) to obtain individualized cells and transferred to a 50mL Falcon tube containing 20% Fetal Bovine Serum (FBS, Gibco) in HBSS to inactivate the papain after the enzymatic digestion. Next, the cell suspension was centrifuged at 200 g for 5 minutes and the resulting pellet was resuspended in 1 mL of DMEM (Gibco) supplemented with 10% FBS and 1% antibiotic-antimycotic (Gibco). Cells were seeded in poly-L-Lysine (15μL/mL, Sigma) coated T-75 flasks with a density of two brains per flask. The medium was changed the day after the culture and every 3-4 days, enriched with the granulocyte-monocyte colony stimulating factor (5ng/mL GM-CSF, Sigma) to promote microglia proliferation at 37°C, 5% CO_2_. When confluence was reached, after approximately 11-14 days, microglia cells were harvested by shaking at 120-140 rpm, at 37°C for 4 hours. The isolated cells were counted and plated at the desired density; 100.000 cells/well in 24- well plates for immunofluorescence experiments, 500.000 cells/well in glass bottom dishes (Ibidi) for live imaging and 2.000.000 cells/well in 6-well plates for Western blot purposes; all coated with poly-L-Lysine to guarantee optimal cell adhesion. Before performing any experiment, microglia were allowed to settle for at least 24 hours.

### BV2 cell line

BV2 cells (Interlab Cell Line Collection San Martino-Instituto Scientifico Tumori-Instituto Nazionale per la Ricerca sul Cancro), a cell line derived from raf/myc-immortalized rat neonatal microglia was used in autophagy assessing experiments wherein transfection of eEGFP-mRFP-LC3 plasmid was required as well as some Western Blot experiments. Cells were grown as an adherent culture in non-coated 90mm^2^ Petri dishes in the presence of 10 mL of DMEM (Gibco) supplemented with 10% FBS and 1% antibiotic-antimycotic (Gibco). When confluence was reached, cells were trypsinized (Trypsin-EDTA 0,5% no phenol red, Gibco) and replated at a 1:5 density. Before performing any experiment, BV2 cells were allowed to settle for at least 24 hours.

### Red fluorescent SH-SY5Y cell line (vampire SH-SY5Y)

The SH-SY5Y cell line is a human neuroblastoma cell line developed as a stable transfection of the SH-SY5Y cell line with the red fluorophore tFP602 (InnoProt, P20303). It derives from neuroepithelioma cell line SK-N-SH, generated from the bone marrow of 4-year-old female with metastatic neuroblastoma. Cells were grown as adherent culture in non-coated T-75 flasks in the presence of 10-15 mL of DMEM (Gibco) supplemented with 10% FBS and 1% antibiotic-antimycotic (Gibco) and 0.25mg/ml Geneticin (G418, Gibco) to select the transfected cells. When confluence was reached, cells were trypsinized (Trypsin-EDTA 0,5% no phenol red, Gibco) and replated at a 1:3 density. SH-SY5Y cells were used to perform phagocytosis assay for which they were trypsinized and replated at the same density 24 hours prior to the phagocytosis assay to avoid the addition of cell clusters to the microglia culture.

### Organotypic hippocampal slice cultures

Organotypic hippocampal slice cultures were prepared as described previously ^67^. Briefly, P7 fms-EGFP pups were sacrificed and brains removed and placed in cold Hank’s Balanced Salt Solution (HBSS) (ThermoFisher Scientific). The hippocampi were extracted and cut into 350 µm slices with a tissue chopper (McIlwain). Slices were then placed in 0.4 µm culture plate inserts (Millipore), each containing 3-5 sections. These inserts were placed in 6 well culture plates, each well containing 1 ml of fresh organotypic slice medium. The medium consisted of 50% Neurobasal medium (Gibco) supplemented with 0.5% B27 (Gibco), 25% horse serum (PAN Biotech), 1% Glutamax (Gibco), 1% antibiotics (penicillin/streptomycin/amphotericin B) (Gibco), and 1% glucose solution in HBSS. The medium was changed the first day after culturing the slices and every 2 days afterwards for 7 days. The experiments were performed on day in vitro 7. For OND, slices were rinsed twice with PBS to remove medium remainders, nutrient deprived in salt solution, and transferred to a hypoxia chamber (1% O_2_) for 3 or 6 hours. For reperfusion experiments, slices were placed back in fresh complete medium containing propidium iodide (PI) (5 µg/ml, Sigma) for an additional hour in a regular cell culture incubator (20% O_2_, 5% CO_2_). The autophagy inhibitor MRT68921 (30 and 100 µM, Sigma) was added to the culture media during the 3hchallenge with OND. The autophagy inducer rapamycin (200 nM, Selleckchem) was tested at 6 and 24h time points and was added for 3 or 21h before the 3h challenge with OND. Immediately after finishing the experiments, the slices were rinsed with PBS and fixed in cold 4% paraformaldehyde solution for 40 minutes. After a couple of rinses with PBS, slices were stored at 4°C until immunofluorescence processing.

### Transient Middle Cerebral Artery occlusion (tMCAo) and rapamycin treatment

2 mo fms-EGFP mice were anaesthetized with isofluorane (2% for induction and 1% for maintenance). tMCAo was induced using the intraluminal filament method as previously described ^68^. Briefly, a midline-neck incision was performed to expose the left common carotid artery (CCA). The left external and internal carotid arteries (ECA and ICA, respectively) were carefully isolated, and a monofilament (silicon coated silk with an extended tip of width of 9-10mm and 0.21mm diameter, Doccol Corporation) was inserted into the ECA and advancing it via the ICA to the origin of the MCA. After a 60min embolization, the monofilament was removed and the CCA reperfused to restore blood flow. The successful occlusion and reperfusion were confirmed by a Laser-Doppler flowmetry (LDF) with a cerebral blood flow (CBF) less than 70%. Mice that did not demonstrate a rapid restoration of the CBF signal during reperfusion were excluded. Body temperature was kept at 37±0.5°C throughout the experiment by an electrical blanket. Mice were sacrificed 6h and 1d after the damage. Sham animals were anesthetized but did not suffer arteries ligation. Some mice received an intraperitoneal injection of rapamycin (10mg/kg, in 20μl/10gr body weight) or vehicle (5% DMSO, 5% Tween-80, 15% polyethylene glycol 400 (PEG400) in 0.9% NaCl solution) during 3 consecutive days. Two injections were administered as a pre-treatment prior to tMCAo surgery and a third injection was given post-tMCAo. Mice were sacrificed 6 hours (h) later.

### Pimonidazole treatment

Mice received one intravascular tail injection of pimonidazole (60mg/kg, diluted in saline) 30min before MCA reperfusion and were sacrificed 30min later. A second set of animals were injected with pimonidazole 30min before sacrifice at 6h and 1d after tMCAo. The percentage of hypoxic area was estimated in confocal tiled images of coronal slices from sham and tMCAo-treated animals. First, hypoxic areas were manually selected in each image using the “Threshold” tool (Fiji) to mask only the pixels of the image with pimonidazole staining. Then, the percentage of pixels occupied by pimonidazole was calculated using the Area Fraction parameter of the “Measure” tool (Fiji). All commands were automated in an ImageJ macro (Fiji). The average area fraction of 4-5 tiled images per animal was calculated.

### Hypoxia-ischemia (HI)

3mo CX3CR1^GFP/+^ CCR2 ^RFP/+^ mice were anesthetized with isoflurane (5% for induction and 2% for maintenance) and the right common carotid artery was ligated with a 6-0 silk suture according to the Rice-Vannucci model ^19, 69^. After ligation, the animals were returned to their home cage for 1h, and then placed in a chamber perfused with a humidified gas mixture (10% oxygen in nitrogen) for 75min at 36°C. Animals were sacrificed 1d and 3d after injury. Control mice were neither subjected to ligation nor hypoxia.

### Cranial irradiation (CIR)

3 mo CX3CR1^GFP/+^ CCR2 ^RFP/+^ mice were irradiated in X-ray irradiator ^20^. Animals were anesthetized with isofluorane (5% for induction and 2% for maintenance) and the whole brain was irradiated with a single dose of 8Gy (0.72Gy/min). The acute exposure of 8Gy is equivalent to approximately 18Gy when delivered in repeated 2Gy fractions, which represents a clinically relevant dose used in treatment protocols such as the medulloblastoma study (PNET5 MB) designed for children (https://clinicaltrials.gov/ct2/show/NCT02066220). Animals were sacrificed 6h and 1d after irradiation. Sham mice were anesthetized but did not receive any CIR.

### Oxygen and nutrient deprivation assay

Primary microglia cultures and hippocampal organotypic slices were treated with oxygen and nutrient deprivation to mimic a hypoxia-like state. Cultures were treated with a salt solution (130 mM NaCl, 5,4 mM KCl, 1.8 mM CaCl_2,_ 26 mM NaHCO_3_, 0.8 mM MgCl_2_, 1.18 mM NaH_2_PO_4_, pH 7.4 in milliQ water) ^70^ without glucose nor amino acids. After adding the salt solution, cultures were placed in a hypoxia chamber (Biospherix, US) inside a standard thermal incubator for 3 or 6 hours with an oxygen concentration between 1-3% and no CO_2_.

### In vitro rapamycin, MRT68921 and bafilomycin-A1 treatment

The autophagy inducer Rapamycin (Selleckchem) was dissolved in DMSO (3.65mM) and diluted in water to 10µM. Rapamycin was added to hippocampal organotypic cultures at a final concentration of 200 nM for 6 and 24 hours. In primary microglia and BV2 cells rapamycin was used at a final concentration of 100 nM for 24 hours and 6 hours, respectively. The autophagy inhibitor MRT68921 (Sigma) was dissolved in water (1 mM). Hippocampal organotypic slices were treated with MRT68921 for 3 hours at a final concentration of 30 and 100 µM. Primary microglia were treated with MRT68921 for 3 and 6 hours at 1, 10, and 30 µM concentrations. The lysosomal inhibitor bafilomycin-A1 (Selleckchem) was dissolved in DMSO (500 µM) and added to primary microglial and BV2 cell cultures at 100 nM concentration for 3h.

### In vitro phagocytosis assay

The *in vitro* phagocytosis assay was adapted from ^67^. Phagocytosis was performed in high glucose DMEM, supplemented with 1% antibiotic-antimycotic (Gibco) and 10% fetal bovine serum (FBS, GE Healthcare Hyclone). Primary microglia were plated and allowed to rest 24 hours prior to the experiment. Cells were fed with apoptotic vampire SH-SY5Y, previously treated with staurosporine (3μM, 4h, Sigma) to induce apoptosis. To ensure collecting only apoptotic cells, the floating fraction was added to primary microglia in a proportion of 1:1 except in the engulfment ratio assay where apoptotic cells were added in a 1:9 proportion. Microglia were left to interact with the apoptotic cells for 1 hour and then fixed to assess the engulfment of the dead cells, or, instead, the non-phagocytosed apoptotic cells were removed by washing twice with PBS and primary microglia left to degrade the engulfed cells for 3 more hours before fixation to assess degradation.

### Two-photon imaging on hippocampal organotypic slices

Organotypic hippocampal slices were obtained from P7 CX3CR1^GFP/+^ mice. Briefly, slices were placed in a fluidic chamber under a Femto-2D microscope equipped with a DeepSee module and GaAsP and PMT detectors. Control slices were perfused with oxygenated complete culture medium whereas OND pre-treated (3h) slices were constantly perfused with nitrogenated OND medium throughout the imaging. The laser was tuned at 920nm to image GFP with a 20X water immersion lens (1.00 N.A.; Olympus). Stacks of 22µm were taken every 1 µm acquired every 60s intervals for 10 minutes (10 timeframes) with a 512 x 512 resolution (0.434µm of pixel width and height).

### Lysosomal pH measurement assay (live imaging)

The measurement of lysosomal pH is based on the ratiometric measurement of two fluorophores by confocal microscopy ^71^: fluorescein (FITC) and tetramethylrohadamine (TRITC) conjugated to a dextran molecule (70,000 MW, anionic, Fisher). Microglia were incubated with 2mg/mL of dextran for 15 hours for its internalization. After that time, the excess of dextran was removed by thorough washing with PBS and the cells incubated in complete medium high glucose DMEM, supplemented with 1% antibiotic-antimycotic (Gibco) and 10% fetal bovine serum (FBS, GE Healthcare Hyclone) for 3.5 hours allowing the dextran to be delivered to the lysosomes (chase pulse). For the OND group the chase was partially done under OND conditions (3h). Next, control cells were washed with imaging medium (150 mM NaCl, 20 mM HEPES, pH 7.4, 1 mM CaCl2, 5 mM KCl, and 1 mM MgCl2, 0.2% glucose) and the OND group was imaged in OND buffer to avoid the loss of the treatment effect. Single plane images were taken using a Leica TCS STED CW SP8 laser scanning microscope using the 63X oil-immersion objective and 2X zoom. Images were analyzed using the Fiji/ImageJ free software and fluorescence intensity was measured for each individual cell. In each experiment, calibration curves were generated from fixed and equilibrated dextranloaded-cells to a range of pH buffers. Calibration groups were fixed in 4% paraformaldehyde for 10 minutes at RT and equilibrated for 20 minutes in the corresponding pH buffer. Buffers contained 50mM Tris Maleate adjusted to 3.5, 4.0, 4.5 and 5.5 pH. Data are shown normalized to the control of each experiment to reduce interexperimental variability.

### Lysosomal enzymatic activity assay

Lysosomal activity was measured using the commercial Lysosomal Intracellular Activity Assay Kit according to the manufactureŕs instructions (Abcam, ab234622). Cells were incubated with a self-quenched substrate, washed with 1mL of ice-cold 1X assay buffer and immediately imaged. Single plane images were taken using a Leica TCS STED CW SP8 laser scanning microscope using the 63X oil-immersion objective and 1.5X zoom. Images were analyzed with Fiji/ImageJ and mean fluorescence intensity was measured for each individual lysosome. Lysosomal fluorescence intensity values proportionally correlate with the amount of lysosomal activity ^72^. Intensity, lysosomal number and occupied area values were normalized to the cell area. Data are shown normalized to the control of each experiment to reduce interexperimental variability

### EGFP-mRFP-LC3 transfection in BV2 cells

The EGFP-mRFP-LC3 plasmid (addgene, #21074) was transfected in Escherichia Coli DH5α competent cells (Invitrogen) using the heat shock method and Kanamycin (50 µg/ml) was used for the selection of transformed bacteria. The plasmid was purified using a maxiprep commercial kit (PureLink HiPure Plasmid Filter Maxiprep Kit, Invitrogen). BV2 cells were seeded in 24 well-plates at 70-90% confluence 24 hours prior to the experiment in high glucose DMEM (Gibco) supplemented with 10% FBS without antibiotics. Transfection assay was performed following manufactureŕs instructions. Lipofectamine 2000 (2µg/mL, Invitrogen) and plasmidic DNA (0.9µg/µL) in a 1:5 ratio were separately pre-incubated with Opti-MEM (Gibco). Lipofectamine 2000 and plasmidic DNA were left to complex for 20 minutes at room temperature and added to the plated BV2 cells in low glucose DMEM supplemented with 5% fetal bovine serum (FBS, GE Healthcare Hyclone) and no antibiotics to ensure full transfection efficiency. Lipofectamine 2000- plasmid complexes were added to BV2 cells for 6 hours and then replaced with the regular growth medium, high glucose DMEM supplemented with 10% fetal bovine serum (FBS, GE Healthcare Hyclone) and 1% antibiotic-antimycotic (Gibco). EGFP-mRFP-LC3 expression was allowed for 24h after the transfection. Cells were fixed in 4% PFA and imaged in a Zeiss LSM 880 Fast Airy Scan microscope under a 40X objective and 2x electronic zoom. A single z-plane was analyzed in Fiji/ImageJ. First, individual puncta were identified using the Find Maxima plugin, after manually optimizing the following parameters: minimum Sigma=2.5, maximum Sigma=3, threshold method Moments, prominence=40 and minimum size=0.032mm. In these puncta, we calculated the GFP/RFP fluorescence ratio as a measure of the degradation of autophagosomes due to the sensitivity of GFP to the lysosomal pH. In addition, we calculated the mean RFP and GFP fluorescence intensity, the total number of puncta, and the area occupied by the puncta per cell and normalized to control conditions (expressed as % change over control).

### Western blot

Primary microglia and BV2 cells were lysed in RIPA buffer (Sigma) containing protease and phosphatase inhibitor cocktail (Fisher). The cell lysate was collected and centrifuged (10.000xg, 10 min). The solubilized protein was quantified by BCA Assay Kit (Fisher) in triplicates at 590nm using a microplate reader (Synergy HT, BioTek). 15 to 20 µg of ß-mercaptoethanol denatured protein were loaded in Tris-glycine polyacrylamide gels (14%) and run for 90 minutes at 120V. The resolved proteins were then transferred to a nitrocellulose membrane at 220mA for 2 hours and the transfer efficiency was verified by Pounceau staining (Sigma). The membranes were then blocked for 1hour in 5% milk prepared in Tris Buffered Saline with 0.1% Tween-20 (TBS-T) buffer. Membranes were afterwards incubated with rabbit primary antibody to LC3 (1:3.000, NB100-2220, Novus Biologicals), and mouse primary antibody to β-actin (1:5.000, Sigma), in TBS-T containing 4% Bovine Serum Albumin (BSA) overnight (4°C, shaker). The next day, membranes were rinsed and incubated with the fluorescent secondary antibodies StarBright Blue 700 anti-mouse (1:5.000, BioRad) and StarBright Blue 700 anti-rabbit (1:5.000, BioRad) in TBS-T containing 5% milk powder. After rinsing membranes, protein bans were imaged in a ChemiDoc imaging system (BioRad). Band intensity was quantified using the Gel Analyzer method of Fiji software.

### Immunofluorescence

Mice were transcardially perfused with 30ml of PBS followed by 30ml of 4% PFA. The brains were removed and post-fixed with the same fixative for 3h at RT, then washed in PBS and kept in cryoprotectant (30% sucrose, 30% ethyleneglycol) at -20°C. Six series of 50µm-thick brain sections were cut using a Leica VT 1200S vibrating blade microtome (Leica Microsystems GmbH, Wetzlar, Germany). Brains obtained from the Karolinska Institute were processed slightly differently. HI-treated mice were anesthetized with 50mg/kg sodium pentobarbital and transcardially perfused-fixed with 6% formaldehyde solution. The tissue was immersion-fixed in the same fixative for 24h at 4°C after perfusion and then soaked overnight in graded concentrations of sucrose solution (10%, 20%, and 30%). The right hemisphere was cut into 40µm-thick sagittal sections in a series of 10 using a sliding microtome (Leica SM2010R, Wetzlar, Germany). The sections were stored in a cryoprotection solution at -20°C for further use. CIR mice were anesthetized with 50mg/kg sodium pentobarbital and transcardially perfused with 4% PFA. The brains were immersion-fixed in the same fixative for 3d at 4°C after perfusion and then soaked overnight in graded concentrations of sucrose solution (10%, 20%, and 30% in 0.1M phosphate buffer). One of the hemispheres was cut into 12 series of 25µm-thick sagittal sections using a sliding microtome (Leica SM2010R, Wetzlar, Germany). The sections were stored in a cryoprotection solution (25% ethylene glycol and 25% glycerin in 0.1M PB) at -20°C for further use. Common marmosets (*Callithrix jacchus*) were deeply anesthetized with isoflurane (5% during 10min) and transcardially perfused with a heparinized solution of saline followed by a solution of 4% PFA. The brains were removed from the skull and post-fixed for 4h in the same fixative. Then the brains were cut in the coronal plane and sections (50μm) were used for staining. The sections were stored in a cryoprotection solution at -20°C for further use. Cynomolgus macaque were deeply anesthetized before lethal injection. The brain was perfused with a saline solution through bilateral carotid catheters, removed from the skull, cut in the coronal plane, post-fixed in a 4% PFA solution and stored at 4°C. It was then soaked in graded concentrations of sucrose solutions in 0.1M PBS (15% for 24 hours followed by 30% for 24 hours) and cut. 40µm sections were stored at -20°C for further use.

Free-floating sections from mice and non-human primates were incubated in permeabilization solution (0.3% Triton-X100, 0.5% BSA in PBS) for 2hr at RT, and then incubated overnight with the primary antibodies diluted in the permeabilization solution at 4°C (a list of primary and secondary antibodies can be found in **Table 1**). Hippocampal organotypic slices were processed similarly, using 0.2% Triton-X-100, 2% BSA in PBS as permeabilization solution. After thorough washing with PBS the sections were incubated with fluorochrome-conjugated secondary antibodies and DAPI (5mg/ml) diluted in the permeabilization solution for 2h at RT. After washing with PBS, the sections were mounted on glass slides with DakoCytomation Fluorescent Mounting Medium (Agilent).

**Table 1.**
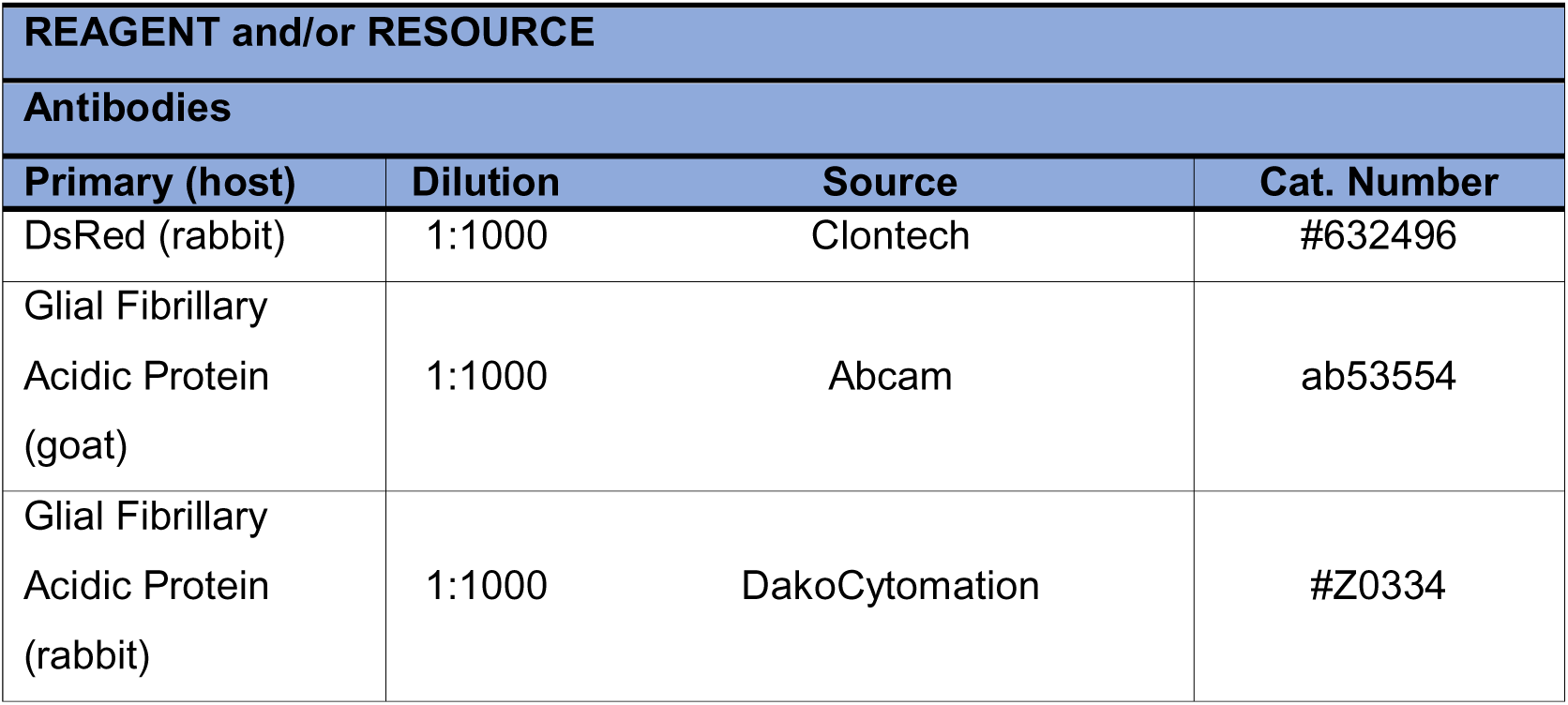
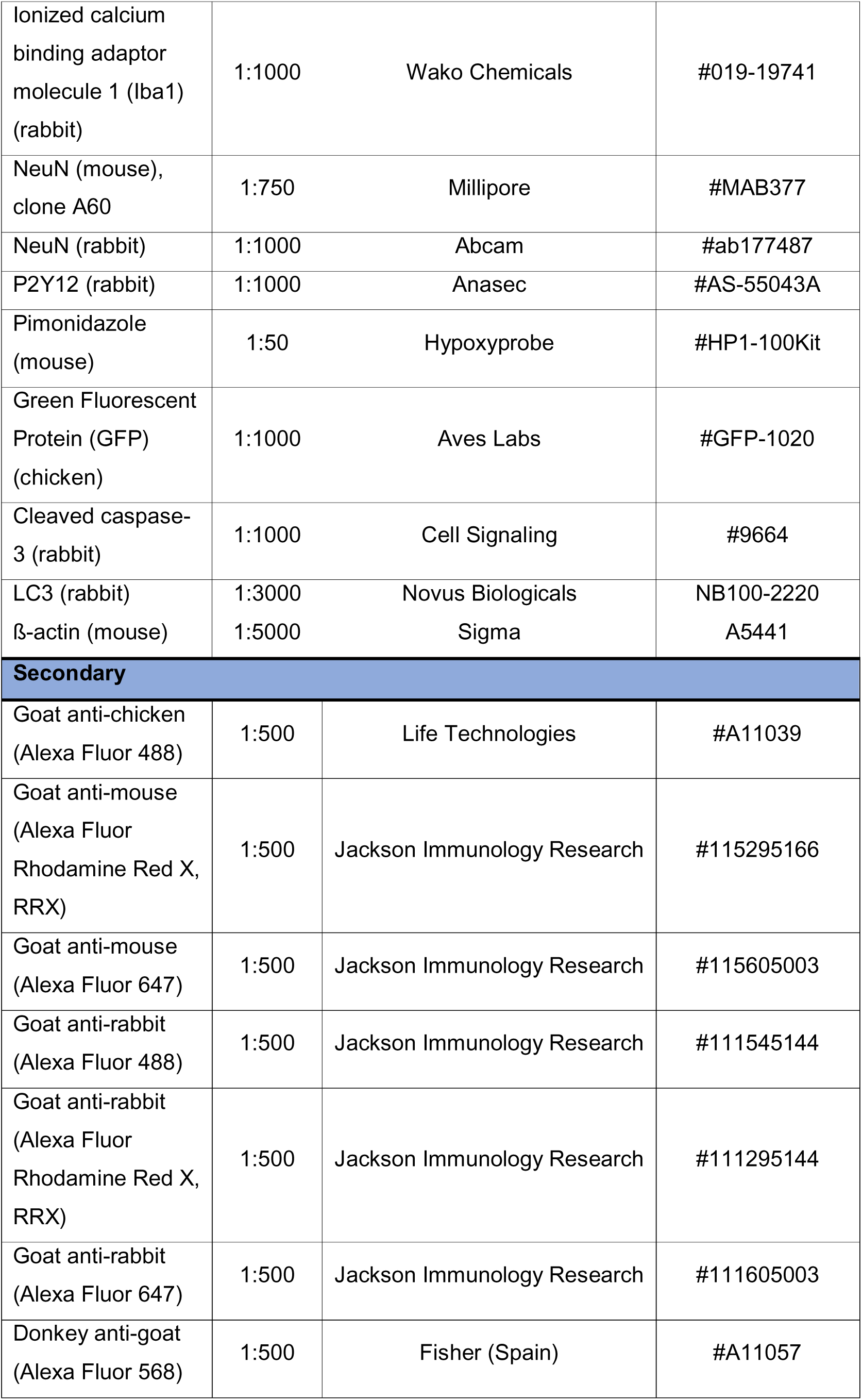
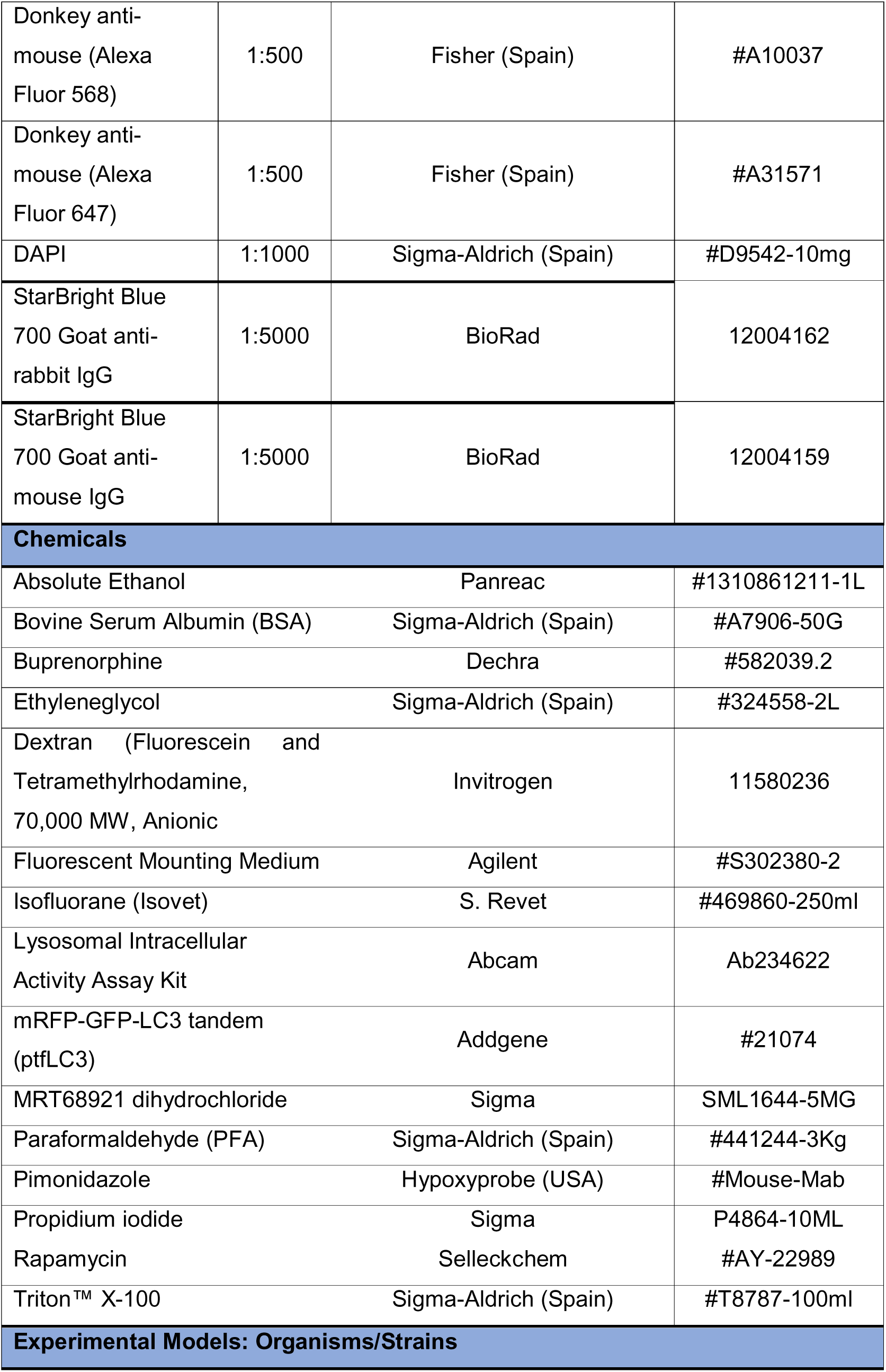
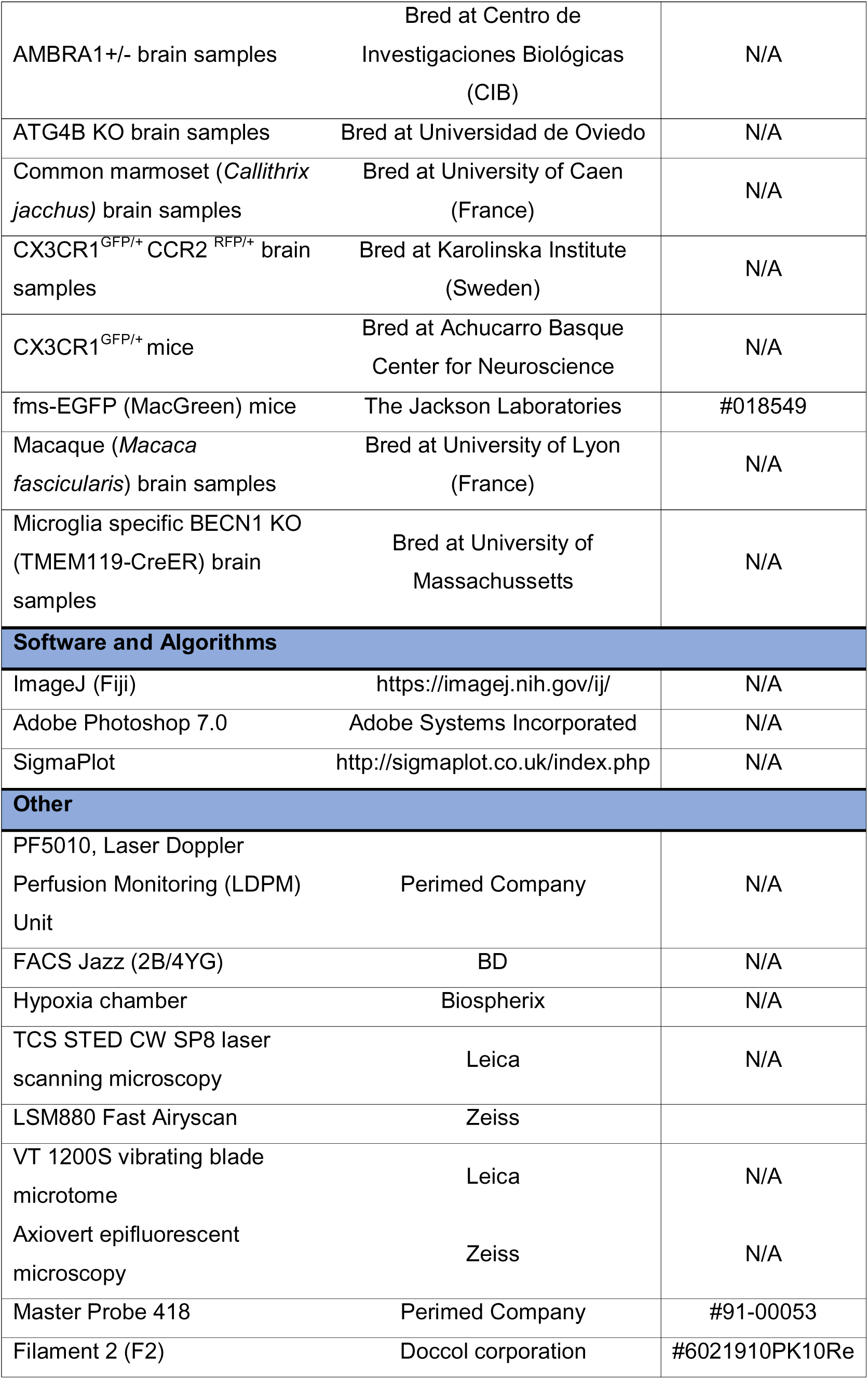
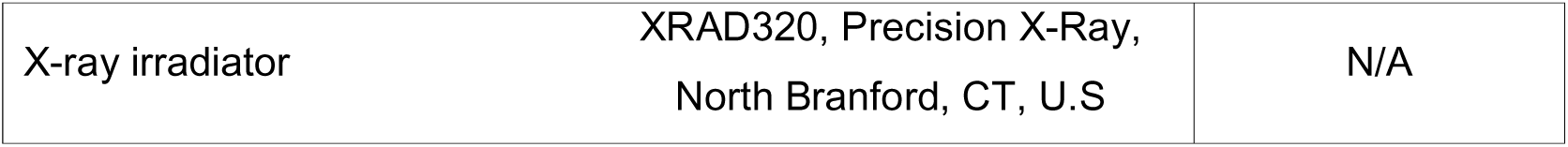
List of reagents and/or resources used in this article.

### Phagocytosis analysis

Fluorescence immunostaining images were collected in a Leica TCS STED CW SP8 laser scanning microscope using 40X or 63X oil-immersion objectives and a z-step of 0.7µm. All images were imported into Adobe Photoshop 7.0 (Adobe Systems Incorporated, San Jose, CA) in tiff format. Brightness, contrast, and background were adjusted equally for the entire image using the “brightness and contrast” and “levels” controls from the “image/adjustment” set of options without any further modification. For mouse tissue sections, 2-5 20µm-thick z-stacks located at random positions containing the DG were collected per hippocampal section, and a minimum of 6 (sagittal) and 4-5 (coronal) sections per series were analyzed. For macaque and common marmoset tissue, one coronal section was fully scanned under the microscope to find all apoptotic cells. In hippocampal organotypic slices, 2-3 images of the DG were acquired per slice and each experimental condition consisted of 3-4 slices. In macaques, the areas covered were cortical regions of the precentral and superior temporal gyri. For primary cultures 3-5 random z-stacks were analyzed from 3 independent coverslips.

Apoptosis and phagocytosis analysis was performed using unbiased stereology methods as previously described ^67^. Apoptotic dead cells were determined by their nuclear morphology visualized with the DNA dye DAPI. These cells lost their chromatin structure (euchromatin and heterochromatin) and appeared condensed and/or fragmented (pyknosis/karyorrhexis). Phagocytosis was defined as the formation of a three dimensional pouch, usually located in the terminal or passant branches of microglia, completely surrounding an apoptotic cell ^14^. In tissue sections and hippocampal organotypic slices the number of apoptotic cells, phagocytosed cells, microglia, and/or monocytes were estimated in the volume of the DG contained in the z-stack (determined by multiplying the thickness of the stack by the area of the DG at the center of the stack using ImageJ (Fiji)). To obtain the absolute numbers (cells per hippocampus), this density value was then multiplied by the volume of the septal hippocampus (spanning from -1mm to -2.5mm in the anteroposterior axes, from Bregma; approximately 6 slices in each of the 6 series), which was calculated using Fiji from images collected at 20x with a Zeiss Axiovert epifluorescent microscope.

In the case of HI and CIR experiments, a limited tissue and/or unrepresentative number of samples was provided by Karolinska Institute (Sweden). Due to this reason, the total volume of the septal hippocampus could not be calculated, and therefore the number of apoptotic, phagocytosed, microglial cells and peripheral monocytes was estimated as cell density (cells per mm^3^).

The following formulae were used to estimate microglial phagocytic efficiency in tissue and organotypic slices:

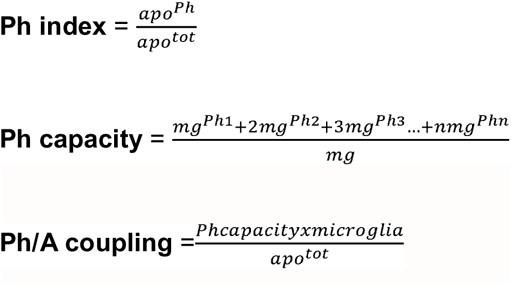

where apo^Ph^ is the number of apoptotic cells phagocytosed; apo^tot^ is the total number of apoptotic cells; and mg^Phn^ is the proportion of microglia with “n” phagocytic pouches.

In primary cultures phagocytosis assays the number of total microglia was calculated, and microglial processes were scanned for DAPI or RFP (from apoptotic SH-SY5Y) inclusions. The number of microglia with inclusions was divided by the total microglial numbers and calculated as percentage of phagocytic microglia at a given timepoint. In the engulfment-degradation experiments, the engulfment was calculated as above, normalized by the control group. The degradation in each experimental condition was calculated by subtracting the percentage of phagocytic microglia at 3h from the percentage of phagocytic microglia in the control group at 1h.

### Transmission Electron Microscopy

Five-million primary microglial cells were cultured in 60 mm diameter plates (Nunclon, ThermoFisher) and exposed to OND in a hypoxia chamber (1% O_2_) with no nutrients (salt solution) during 3 hours. Subsequently, primary microglia were rinsed in PBS and pre-fixed as an adherent cell monolayer using 0.5% glutaraldehyde solution in Sörenson buffer 0.1M pH=7.4 (SB) (10 mins, RT). After scraping the pre-fixed cells, primary microglia were centrifuged (800g, 5 mins, RT) to form a pellet and fixed in 2% glutaraldehyde solution in SB (overnight, 4°C). Primary microglia were then rinsed with 4% sucrose in SB and post-fixed with 1% osmium tetroxide in SB (1h, 4°C, darkness). After rinsing, primary microglia were dehydrated in a growing concentration series of acetone. After dehydration, primary microglia samples were embedded in epoxy resin EPON Polarbed 812 (Electron Microscopy Sciences). Semi-thin sections (1 µm thick) were stained with toluidine blue to identify the regions of interest. Ultra-thin sections were cut using a LEICA EM UC7 ultramicrotome and contrasted with uranyl-acetate and lead citrate. The ultrastructural analysis was done with a Transmission Electron Microscope Jeol JEM 1400 Plus at 100 kVs equipped with a sCMOS digital camera. Image analysis (4-6 images per cell) was performed in 36-38 cells per experimental condition using Fiji. Autophagic-like vesicles (containing at least a portion of double membrane with granular, membranous and heterogeneous cargo) and lysosomal-like vesicles (electron-dense vesicles with single or double membrane) were identified manually and their perimeter was selected to generate regions of interest (ROIs) in each image. The area of the cellular cytoplasm (µm^2^) was also identified and selected manually to generate a ROI in each image. Subsequently, the number (vesicles) and area (vesicles and cytoplasm, µm^2^) of ROIs were measured. Finally, the quantitative data coming from ROIs of the images belonging to the same cell were grouped and the number of autophagic-and lysosomal-like vesicles per µm^2^ was calculated dividing the number of vesicles present in the cell by its cytoplasm area (µm^2^). The percentage of cytoplasm area occupied by autophagic- and lysosomal-like vesicles per each cell was calculated by summing up the areas of individual vesicles and relating it to the cellular cytoplasmic area, which was considered 100%. The mean size (area, µm^2^) of the vesicles per cell is represented in logarithmic scale for illustrative purposes.

### Statistical analysis

SigmaPlot (San Jose, CA, USA) was used for statistical analysis. Data was tested for normality and homoscedasticity (Levenés test). When the data did not comply with these assumptions, a logarithmic transformation (Log_10_, Log_10_+1, of Ln) or a square root was performed and the data was analyzed using parametric tests. Two-sample experiments were analyzed by Studentś t-test and more than two sample experiments with one-way or two-way ANOVA. In two-way ANOVAs, the interaction between factors was assessed prior to analyzing the effect of individual factors. In case that homoscedasticity or normality were not achieved with a logarithmic transformation, data was analyzed using a Kruskal-Wallis ranks test, followed by Dunn method as a *post hoc test.* Two sample non-parametric data was analyzed using Mann Whittney U test. Only p < 0.05 is reported to be significant.

---

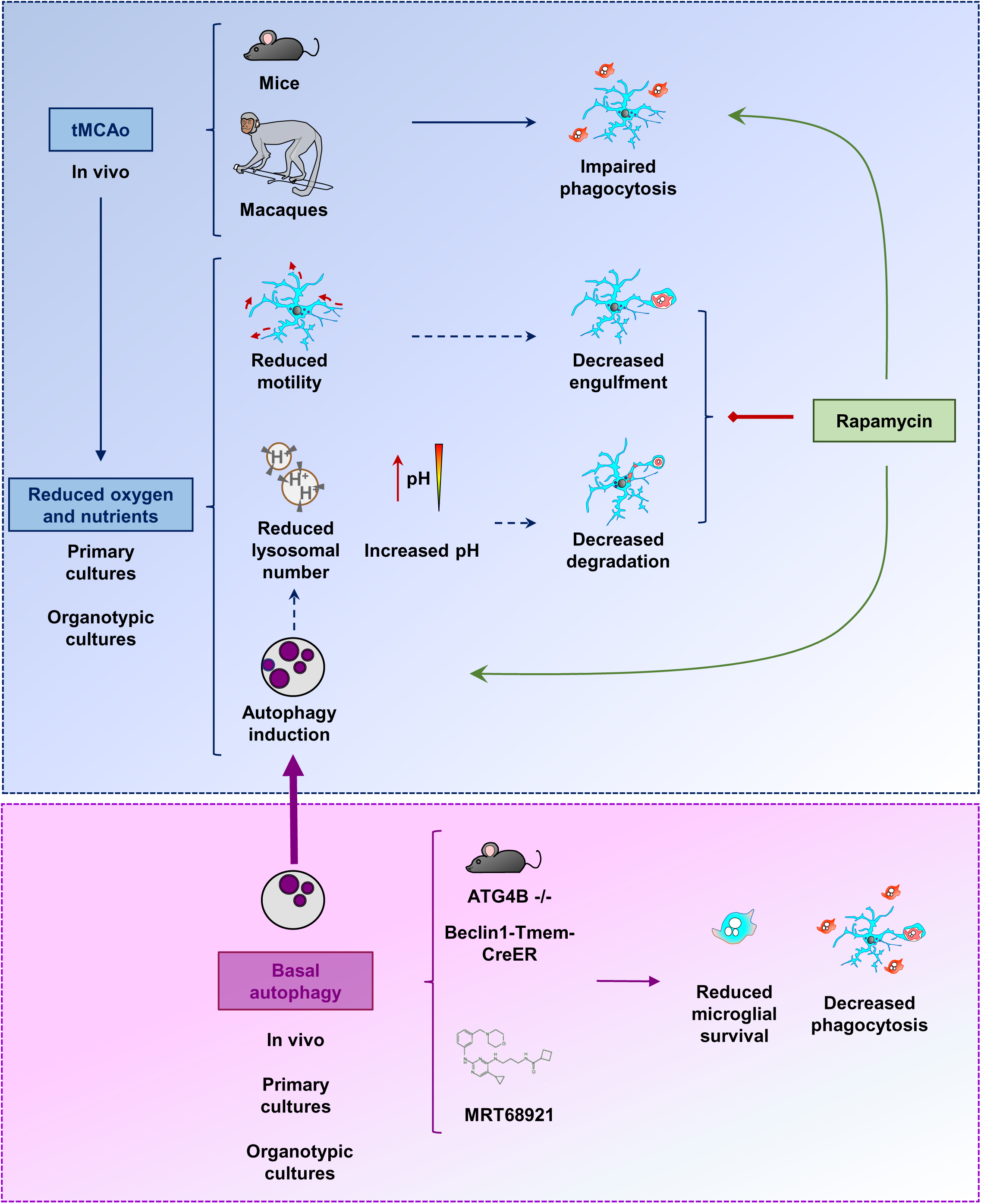

## Notes

### Competing Interest Statement

The authors have declared no competing interest.

